# Efficient parameter calibration and real-time simulation of large scale spiking neural networks with GeNN and NEST

**DOI:** 10.1101/2022.05.13.491646

**Authors:** Felix Johannes Schmitt, Vahid Rostami, Martin Paul Nawrot

## Abstract

Spiking neural networks (SNN) represent the state-of-the-art approach to the biologically realistic modeling of nervous system function. The systematic calibration for multiple free model parameters is necessary to achieve robust network function and demands high computing power and large memory resources. Special requirements arise from closed-loop model simulation in virtual environments, and from real-time simulation in robotic application. Here, we compare two complementary approaches to efficient large scale and realtime SNN simulation. The widely used NEural Simulation Tool (NEST) parallelizes simulation across multiple CPU cores. The GPU-enhanced Neural Network (GeNN) simulator uses the highly parallel GPU-based architecture to gain simulation speed. We quantify fixed and variable simulation costs on single machines with different hardware configurations. As benchmark model we use a spiking cortical attractor network with a topology of densely connected excitatory and inhibitory neuron clusters with homogeneous or distributed synaptic time constants and in comparison to the random balanced network. We show that simulation time scales linearly with the simulated biological model time and, for large networks, approximately linearly with the model size as dominated by the number of synaptic connections. Additional fixed costs with GeNN are almost independent of model size, while fixed costs with NEST increase linearly with model size. We demonstrate how GeNN can be used for simulating networks with up to 3.5 · 10^6^ neurons (> 3 · 10^12^ synapses) on a high-end GPU, and up to 250, 000 neurons (25 ·10^9^ synapses) on a low-cost GPU. Real-time simulation was achieved for networks with 100, 000 neurons. Network calibration and parameter grid search can be efficiently achieved using batch processing. We discuss the advantages and disadvantages of both approaches for different use cases.

## Introduction

Information processing in animal nervous systems is highly efficient and robust. The vast majority of nerve cells in invertebrates and vertebrates are action potential generating (aka spiking) neurons. It is thus widely accepted that neural computation with action potentials in recurrent networks forms the basis for sensory processing, sensory-to-motor transformations and higher brain function (Abeles, 1991; Singer and Gray, 1995). The availability of increasingly detailed anatomical, morphological and physiological data allows for well-defined functional SNNs of increasing complexity that are able to generate testable experimental predictions at physiological and behavioral levels. SNNs have thus become a frequent tool in basic (Brunel, 2000; Van Vreeswijk and Sompolinsky, 1996), translational (McIntyre and Hahn, 2010; Eliasmith et al., 2012), and clinical (Kasabov and Capecci, 2015; Hammond et al., 2007) neuroscience research. In the applied sciences, brain-inspired SNNs have the potential to shape future solutions for intelligent systems (Neftci et al., 2013; Chicca et al., 2014; Schuman et al., 2022). This specifically includes spike-based approaches to machine learning (Indiveri et al., 2010; Schmuker et al., 2014; Zenke and Ganguli, 2018; Gütig and Sompolinsky, 2006; Gütig, 2016; Rapp et al., 2020; Pfeiffer and Pfeil, 2018; Tavanaei et al., 2019), to reservoir computing (Büsing et al., 2010; Tanaka et al., 2019), and brain-inspired control architectures for artificial agents and robots (Feldotto et al., 2022; Sakagiannis et al., 2021; Rapp and Nawrot, 2020; Bartolozzi et al., 2022; Helgadóttir et al., 2013). Computation with attractor networks has been hypothesized as one hallmark of brain-inspired computation (Hopfield, 1982; Amit and Brunel, 1997) and, with increasing evidence, has been implicated in decision-making (Finkelstein et al., 2021), working memory (Sakai and Miyashita, 1991; Inagaki et al., 2019), and sensory-motor transformation (Mazzucato et al., 2019; Wyrick and Mazzucato, 2021; Rostami et al., 2022; Mazzucato, 2022).

The conventional tool for the simulation of SNNs are CPU-based simulation environments. Several well-adopted simulators are in community use (Brette et al., 2007; Tikidji-Hamburyan et al., 2017), each of which has typically been optimized for specific purposes such as the simulation of complex neuron models with extended geometry and detailed biophysics (Hines and Carnevale, 2001), the convenient implementation of neuron dynamics by means of coupled differential equations (Stimberg et al., 2019), or the implementation of the Neural Engineering Framework (NEF) (Eliasmith and Anderson, 2003; Bekolay et al., 2014). The NEural Simulation Tool (NEST, https://www.nest-simulator.org/, (Gewaltig and Diesmann, 2007)) that we consider here was designed for the parallelized simulation of large and densely connected recurrent networks of point neurons. It has been under continuous development since its invention under the name of SYNOD (Diesmann et al., 1995, 1999; Rotter and Diesmann, 1999; Morrison et al., 2005, 2007; Jordan et al., 2018) and enjoys a stable developer and a large user community. More recently, new initiatives have formed to harness GPU-based simulation speed for the modeling of SNNs (Nageswaran et al., 2009; Fidjeland et al., 2009; Mutch et al., 2010; Brette and Goodman, 2012; Florimbi et al., 2021; Golosio et al., 2021; Ben-Shalom et al., 2022). The GPU-enhanced Neural Network (GeNN) simulation environment (https://genn-team.github.io/genn/) developed by Thomas Nowotny and colleagues (Yavuz et al., 2016; Knight et al., 2021; Knight and Nowotny, 2021) is a code generation framework (Blundell et al., 2018) for SNNs and their use in computational neuroscience and for machine learning (Knight and Nowotny, 2022). Neuromorphic hardware (Ivanov et al., 2022; Javanshir et al., 2022) provides an alternative substrate for the simulation of SNNs and is not considered here.

Aim of the present study is to evaluate the use of a GPU-based simulation technique (GeNN) in comparison with a CPU-based simulation technique (NEST) with respect to simulation speed in dependence of network size and in the context of efficient parameter search. We restricted our benchmark approach to simulations on single machines with multiple CPU cores that we consider standard equipment in a computational lab and we compare simulation performance on a high-end GPU with an affordable low-cost GPU that can be used e.g. for teaching purposes.

## Results

### Spiking neural attractor network as benchmark model

We performed simulations of the spiking cortical attractor network model established in Rostami et al. (2022). This network inherits the overall network structure of the random balanced network (RBN, Van Vreeswijk and Sompolinsky 1996; Brunel 2000) with random recurrent connections (drawn from the Bernoulli distribution) among excitatory and inhibitory neurons (Fig. 1A) but introduces a topology of strongly interconnected pairs of excitatory and inhibitory neuron populations (E/I clusters, Fig. 1B) by increasing the intra-cluster synaptic weights (see Material and Methods). This E/I clustered network exhibits a complex pattern of spontaneous network activity, where each cluster can dynamically switch between a state of low (baseline) activity and states of increased activity (Fig. 1D). This network behavior marks the desired feature of metastability (Mazzucato et al., 2019; Rost et al., 2018; Rostami et al., 2022) where the network as a whole cycles through different attractors (or network-wide states) that are defined by the possible cluster activation patterns.

**Figure 1.**
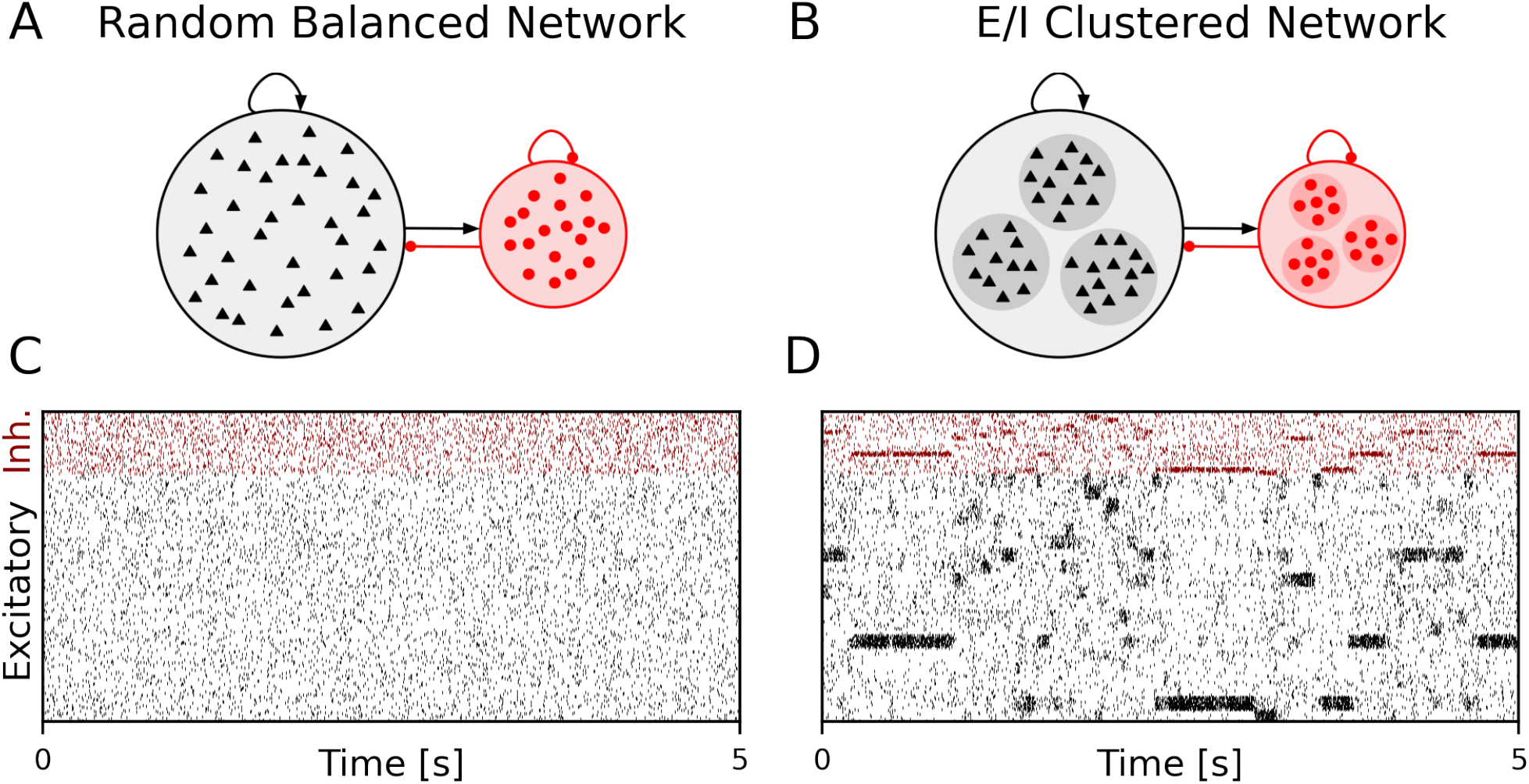
Metastable network activity emerges by introducing excitatory-inhibitory clusters in the random balanced network. (A) Sketch of RBN architecture with one excitatory neuron pool (gray shaded circle, 80% of all neurons) and one inhibitory neuron pool (red shaded circle, 20% of all neurons). Excitatory neurons (black triangles) and inhibitory neurons (red circles) make random connections within and across pools. The respective connection strengths are tuned such that for each neuron, on average, the total synaptic input current balances excitatory and inhibitory input currents. (B) Sketch of excitatory-inhibitory (E/I) cluster topology. Both, excitatory and inhibitory neuron pools are tiled into clusters (small shaded circles) of strongly interconnected neurons (indicated by darker shading). In addition, each excitatory cluster is strongly and reciprocally connected to one corresponding inhibitory cluster such that the balance of excitatory and inhibitory synaptic input is retained for all neurons. In our network definition a single parameter *J*_E+_ determines the cluster strength in terms of synaptic weights and allows to move from the RBN (*J*_E+_ = 1) to increasingly strong clusters by increasing *J*_E+_ > 1. The model uses exponential leaky Integrate-and-Fire (I&F) neurons and all neurons receive weak constant input current. (C-D) Raster plot of excitatory (black) and inhibitory (red) spiking activity in a network of *N* = 25, 000 neurons during 5 s of spontaneous activity after an initial warm-up time of one second was discarded. Shown are 8% of the total neuron population. The spike raster plots are generated from GeNN simulations. (C) The RBN (*J*_E+_ = 1.0, *I*_thE_ = 2.6, *I*_thI_ = 1.9) exhibits irregular spiking of excitatory and inhibitory neurons with constant firing rates that are similar for all excitatory and inhibitory neurons, respectively. (D) The E/I clustered network (*J*_E+_ = 2.75, *I*_thE_ = 1.6, *I*_thI_ = 0.9) shows a metastable behavior, where different individual E/I clusters can spontaneously assume states of high activity and fall back to spontaneous activity levels. The overall firing rates are higher than in the RBN.

The pairwise Bernoulli connectivity scheme with a connection probability *p* between any pair of neurons implies that the number of synapses *M* scales quadratically with the number of neurons *N* as *M* = *pN* ^2^. For the chosen network parameters we obtain an overall connectivity parameter of *p* ≈ 0.3 (see Materials and Methods). The clustered network topology in our benchmark model results from stronger synaptic excitatory and inhibitory weights within each E/I cluster than between different E/I clusters (Fig. 1B). This compartmentalized architecture suits well for our benchmarking purpose because it is reminiscent for whole-system or multi-area modeling in large scale models that involve several neuropiles or brain areas (Rapp and Nawrot, 2020; Schmidt et al., 2018). We kept the number of clusters fixed to *N*_*Q*_ = 20.

### Benchmark approach and quantification of simulation costs

We benchmark performance by measuring the wall-clock time of the simulation. We differentiate fixed costs *T*_fix_ that are independent of the biological model time to be simulated, and variable costs *T*_var_ determined by the simulation speed after model generation (see Materials and Methods). We used two different hardware configurations for CPU-based simulation with NEST (servers S2 and S3 in Table 1) and two hardware configurations for GPU-based simulation with GeNN (Table 1) comparing a low-cost GPU (S1) with a state-of-the-art high-end GPU (S3). With GeNN we tested two different approaches to store the connectivity matrix of the model (Knight and Nowotny, 2021). The `SPARSE` connectivity format (Sp) stores the matrix in a sparse representation. The `PROCEDURAL` connectivity (Pr) regenerates the connectivity on demand, i.e. after a spike has occurred.

**Table 1.** Hardware configurations and benchmark setups.

| CPU |  |  | GPU |  |  |  |
| --- | --- | --- | --- | --- | --- | --- |
| No. of cores | Clock speed [GHz] | Memory [GB] | Architecture | No. of CUDA-cores | Memory [GB] | Performance (single) [TFLOPS] |
| **Server 1 (S1) Ubuntu 16.04 LTS** |  |  |  |  |  |  |
| Dual AMD Opteron 6380 |  |  | GeForce GTX 970 |  |  |  |
| 2 × 16 | 2.5 | 128 | Maxwell | 1,664 | 4 | 3.5 |
| **Server 2 (S2) Ubuntu 16.04 LTS** |  |  |  |  |  |  |
| Dual Intel Xeon E5-2630 v4 |  |  | - |  |  |  |
| 2 × 10 | 2.2 | 192 | - | - | - | - |
| **Server 3 (S3) Ubuntu 20.04 LTS** |  |  |  |  |  |  |
| Intel Xeon Gold 6248R |  |  | Quadro RTX A6000 |  |  |  |
| 24 | 3.0 | 128 | Ampere | 10,752 | 48 | 38.7 |

### Fixed costs for GeNN are high but independent of network size

We find that for NEST the overall fixed costs scale approximately linearly with the network connectivity as expressed in the total number of connections *M* ∝ *N* ^2^ (Fig. 2) while the overall fixed costs stay approximately constant for GeNN and essentially over the complete range of tested network sizes. The fixed costs add up different contributions as shown in Fig. 2A. These are model definition, building of the model, and loading of the model for GeNN and node creation and creation of connections for NEST (see Materials and Methods). For NEST, compilation of the model (Build phase of GeNN) is not needed because it uses pre-compiled neuron and synapse models (Diesmann and Gewaltig, 2002) in combination with exact integration (Rotter and Diesmann, 1999). The fixed costs of GENN are dominated by the wall-clock time required for building the model and these appear to be essentially independent of model size. The costs of model definition and loading the model increases with model size, but make only a negligible contribution to the overall fixed costs. Thus, for a small network size of *N* = 5, 000 neurons the overall fixed costs amount to ≈ 30 s and ≈ 3 min for the `PROCEDURAL` and `SPARSE` connectivity, respectively, compared to only 3 s with NEST. This picture changes for a ten times larger network with *N* = 50, 000 neurons. Now, the wall-clock time for setting up the model with NEST is more costly than building with GeNN (`PROCEDURAL` connectivity) on our hardware configurations. The fixed costs for NEST increase quadratically in *N* (linear in *M*) on both hardware configurations and eventually exceed fixed costs with GeNN (Fig. 2B).

**Figure 2.**
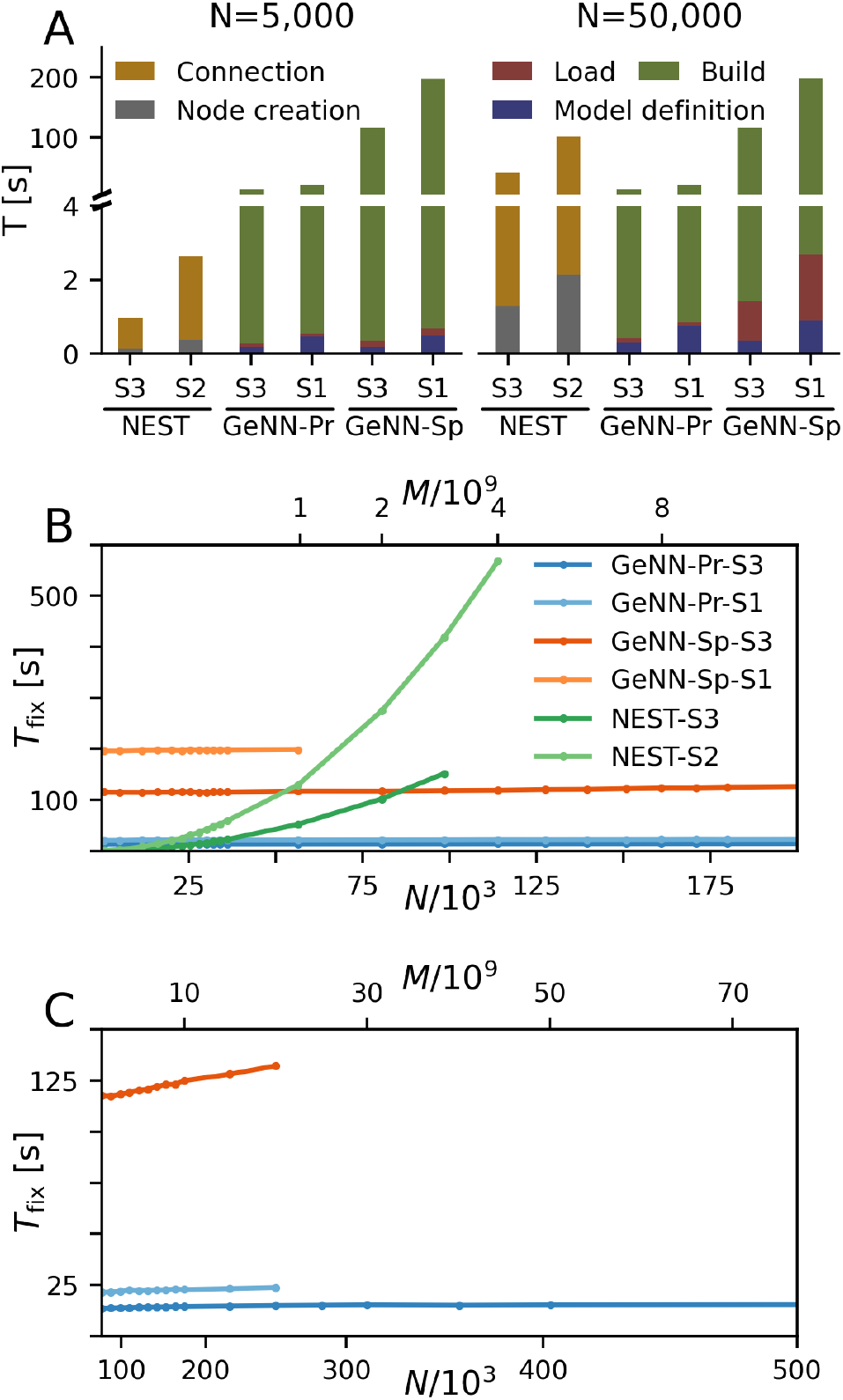
Fixed costs of simulation. (A) Individual costs for execution phases of NEST and GeNN for two network sizes of *N* = 5, 000 and *N* = 50, 000 neurons. The GPU-based simulation requires expensive model compilation (Build). NEST uses pre-compiled neuron models. Note that the y-axis has different linear scales for low (≤4) and high (≥4) values of fixed costs. (B,C) Total fixed costs in seconds over network size for the different simulators and hardware configurations as indicated. Total fixed costs are approximately constant across network size for the GPU-based simulation with the PROCEDURAL connectivity, while they have a small slope for the SPARSE connectivity. Panel C extends panel B for larger network sizes. Note that the x-axis in panel B is linear in *N*, while the x-axis in panel C is linear in *M*.

The maximal network size that we were able to simulate is indicated by the end points in the benchmark graphs in Fig. 2B and C. It is bound by the available memory, which is required for storing the network model and the data recorded during simulation. In the case of simulation with NEST, this bound is determined by the RAM configuration (Table 2). The larger RAM size of 192 GB on S2 allowed for a maximum network connectivity of *M* = 4 ·10^9^ synapses and *N* ≈ 114, 000 neurons, while on the faster server configuration S3 with 128 GB RAM the limit was reached earlier (Fig. 2B). With GeNN the limiting factor for the network size and connectivity is the hardware memory on the GPU itself. The `PROCEDURAL` connectivity allows for a more efficient usage of the GPU memory at the expense of simulation speed and allowed for a network size of > 3.5 ·10^6^ neurons and > 3, 000 ·10^9^ synapses on the high-end GPU (S3) and a respectable size of *N* ≈ 250, 000 neurons (*M* ≈ 20 ·10^9^ synapses) on the low-cost GPU (S1).

**Table 2.**
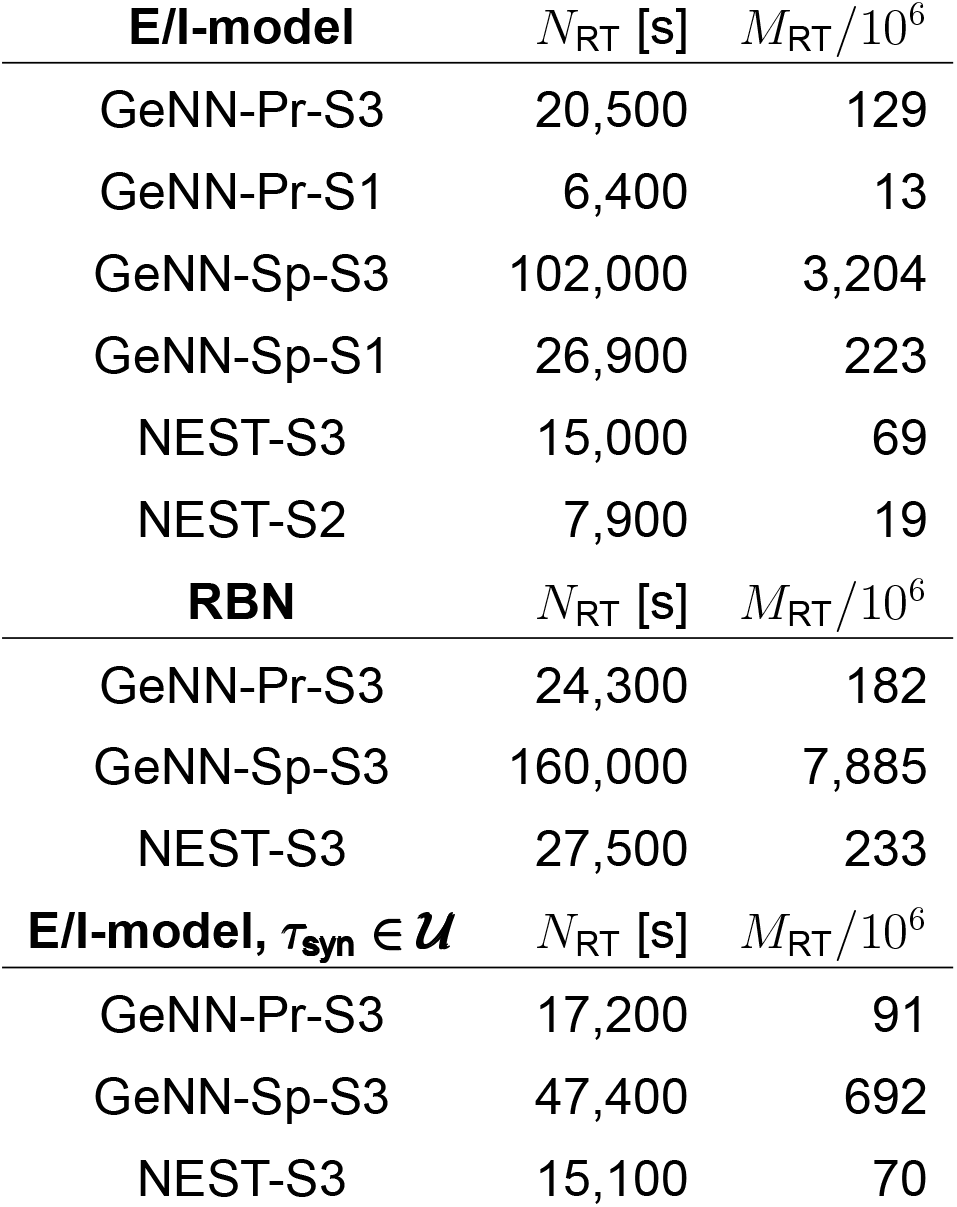
Maximum network size *N*_RT_ within real-time limit. The real-time limit for the E/I-model was determined in simulation steps of 500 neurons. The real-time limits of the RBN and E/I-model with heterogeneous synaptic time constants were estimated with a linear interpolation between the nearest data points (cf. markers in Fig. 4C). Simulations comprised 10 seconds of biological model time with Δ*t* = 0.1 ms.

| **E/I-model** | $N_{\text{RT}}$ [s] | $M_{\text{RT}}/10^6$ |
| --- | --- | --- |
| GeNN-Pr-S3 | 20,500 | 129 |
| GeNN-Pr-S1 | 6,400 | 13 |
| GeNN-Sp-S3 | 102,000 | 3,204 |
| GeNN-Sp-S1 | 26,900 | 223 |
| NEST-S3 | 15,000 | 69 |
| NEST-S2 | 7,900 | 19 |
| **RBN** | $N_{\text{RT}}$ [s] | $M_{\text{RT}}/10^6$ |
| GeNN-Pr-S3 | 24,300 | 182 |
| GeNN-Sp-S3 | 160,000 | 7,885 |
| NEST-S3 | 27,500 | 233 |
| **E/I-model, $\tau_{\text{syn}} \in \mathcal{U}$** | $N_{\text{RT}}$ [s] | $M_{\text{RT}}/10^6$ |
| GeNN-Pr-S3 | 17,200 | 91 |
| GeNN-Sp-S3 | 47,400 | 692 |
| NEST-S3 | 15,100 | 70 |

### Variable costs scale linearly with biological model time and approximate linearly with network connectivity

We first quantify wall-clock time *T*_var_ in dependence on the simulated biological model time *T*_bio_ for a fixed network size of *N* = 50, 000 (Fig. 3A). As to be expected, simulation time grows approximately linearly with the number of simulated time steps and, thus, for a predefined simulation time constant 1/Δ*t*, is proportional to the simulated time with

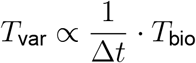

for all tested hardware platforms. We see considerably faster simulations with the `SPARSE` connectivity compared to the `PROCEDURAL` connectivity in our network with a total connectivity of *p* ≈ 0.308. Interestingly, a previous publication by Knight and Nowotny (2021) considering the RBN with a lower connectivity density of *p* = 0.1 found that `PROCEDURAL` connectivity performs equally fast or even faster, which could be due to their lower connectivity density.

**Figure 3.**
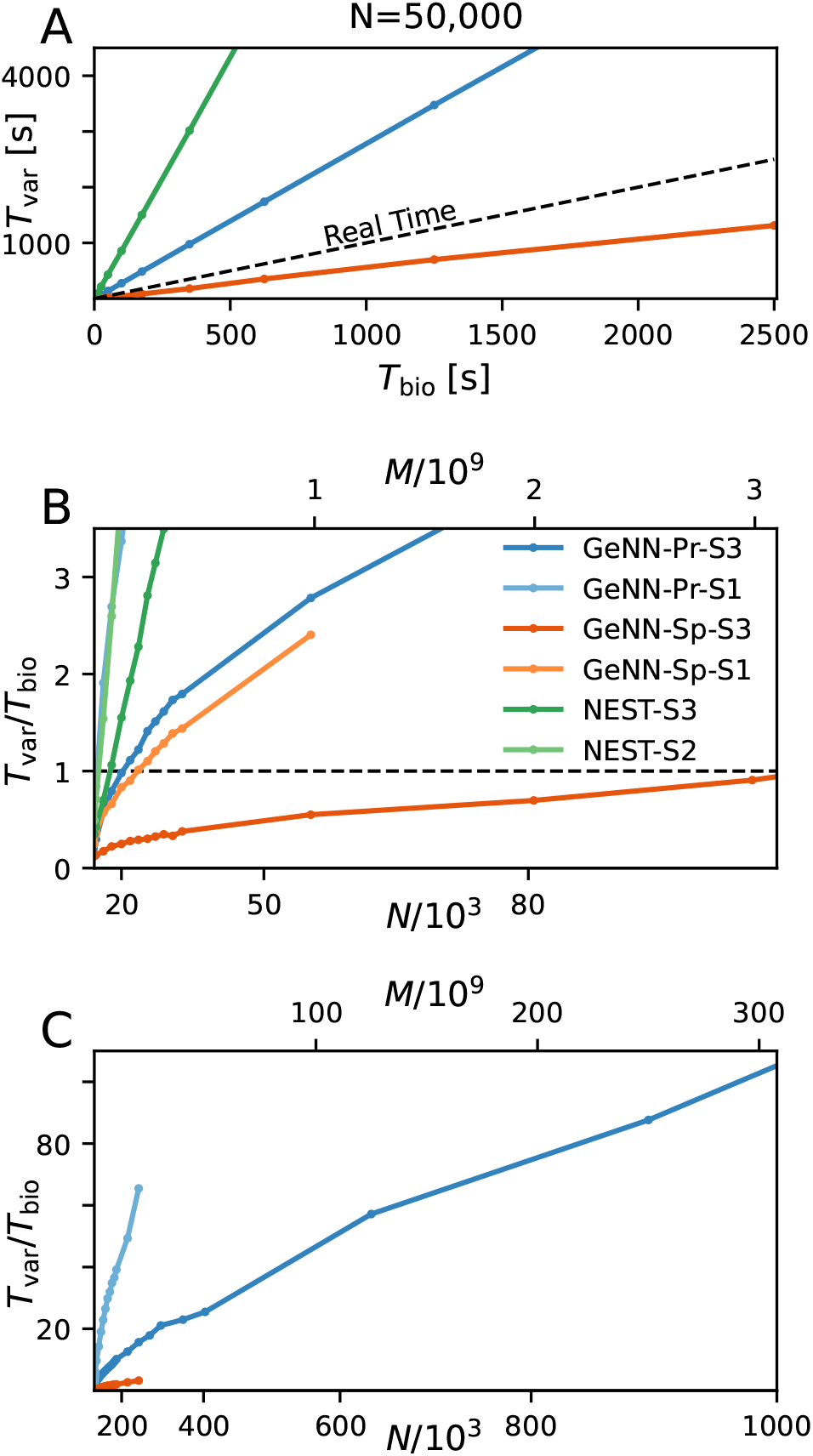
Variable costs of simulation and real-time limitation. (A) Wall-clock time *T*_var_ as a function of simulated biological model time *T*_bio_. (B) Real time factor *RT* = *T*_var_/*T*_bio_ as a function of network size. Real-time capability requires a variable cost factor that remains below *RT* = 1 (dashed line). On the high-end GPU, simulation faster than real-time can be achieved for a network size of *N* ≈ 100, 000 neurons and *M* ≈ 3.1 ∝10^9^ synapses. With the best performing NEST configuration, a network of *N* ≈ 15, 000 neurons and *M* ≈ 0.1 ∝10^9^ synapses can be simulated in real-time. GeNN-Pr-S1 is congruent with NEST-S2 in this view. (C) As in B for larger network size. Wall-clock time scales almost linearly with the total number of synapses *M* (and thus with *N* ^2^) for large network sizes. All simulations have been conducted with a time resolution of Δ*t* = 0.1 ms in biological model time. (B, C) The x-axis is linear in M (top axis).

Next, we analyzed the relation between wall-clock time and network size. As shown in Fig. 3B and C, the the proportionality factor *T*_var_/*T*_bio_ for NEST shows an approximate linear dependence on the total number of synapses *M* for large network sizes and the variable costs are thus proportional to the squared number of neurons *T*_var_ ∝ *pN* ^2^ · *T*_bio_ with linear scale factor *p*, denoting the network-specific connectivity.

For GeNN the graph shows a convex relation between *T*_var_/*T*_bio_ and lower range network sizes. For increasing network size this relation becomes increasingly linear in *N* ^2^. Additional model-related factors may contribute, in particular the average spiking activity of neurons and the resulting spike traffic (see Discussion).

Real-time simulation is defined as *T*_sim_ = *T*_bio_. For For applications in neurorobotics or when using SNNs for real-world machine learning application we may require that the simulation runs equally fast or faster than real-time. For our attractor network model we determined a maximum network size of *N*_RT_ = 102, 000 that fulfills the real-time requirement using GeNN on the high-end GPU (Fig. 3B, Table 2).

The RBN is a standard model in computational neuroscience and widely for the simulation of cortical activity. We therefore repeated our calibrations for the RBN in direct comparison to the E/I clustered network using the fastest hardware configuration (S3). The fixed costs are identical for both network types. This was to be expected as the overall network connectivity is identical in both cases. The variable costs show the same general dependence on network size but are smaller for the RBN mainly due to the overall lower firing rates (Fig.1).

### Simulation costs with heterogeneous synaptic time constants

Thus far all neuron and synapse parameters were identical across the network with fixed synaptic weight and time constant for excitatory and inhibitory synapses, respectively. We now introduce heterogeneity of the excitatory and inhibitory synaptic time constants using uniform distributions with the means corresponding with the parameter values used before and a standard deviation of *±*5%. Using the same neuron model in NEST as above, the heterogeneity applies across postsynaptic neurons while for each neuron all incoming synapses have identical time constants for excitatory and inhibitory synapses, respectively (see Materials and Methods). In GeNN we defined the neuron model and synapse model independently and synaptic time constants are heterogeneously distributed across all excitatory and inhibitory synapses individually, independent of postsynaptic neuron identity (see Discussion).

As a first result we observe that the E/I clustered network retains the desired metastable network dynamics with distributed synaptic time constants as shown in Fig. 4A. When comparing the simulation costs to the homogeneous case on the fastest hardware configurations (S3) we find that fixed costs did not increase with NEST and show an indistinguishable dependence on network size (Fig. 4B). However, with GeNN fixed costs remain independent of network size but were strongly increased in comparison to the homogeneous case, both for the `SPARSE` and the `PROCEDURAL` approach.

**Figure 4.**
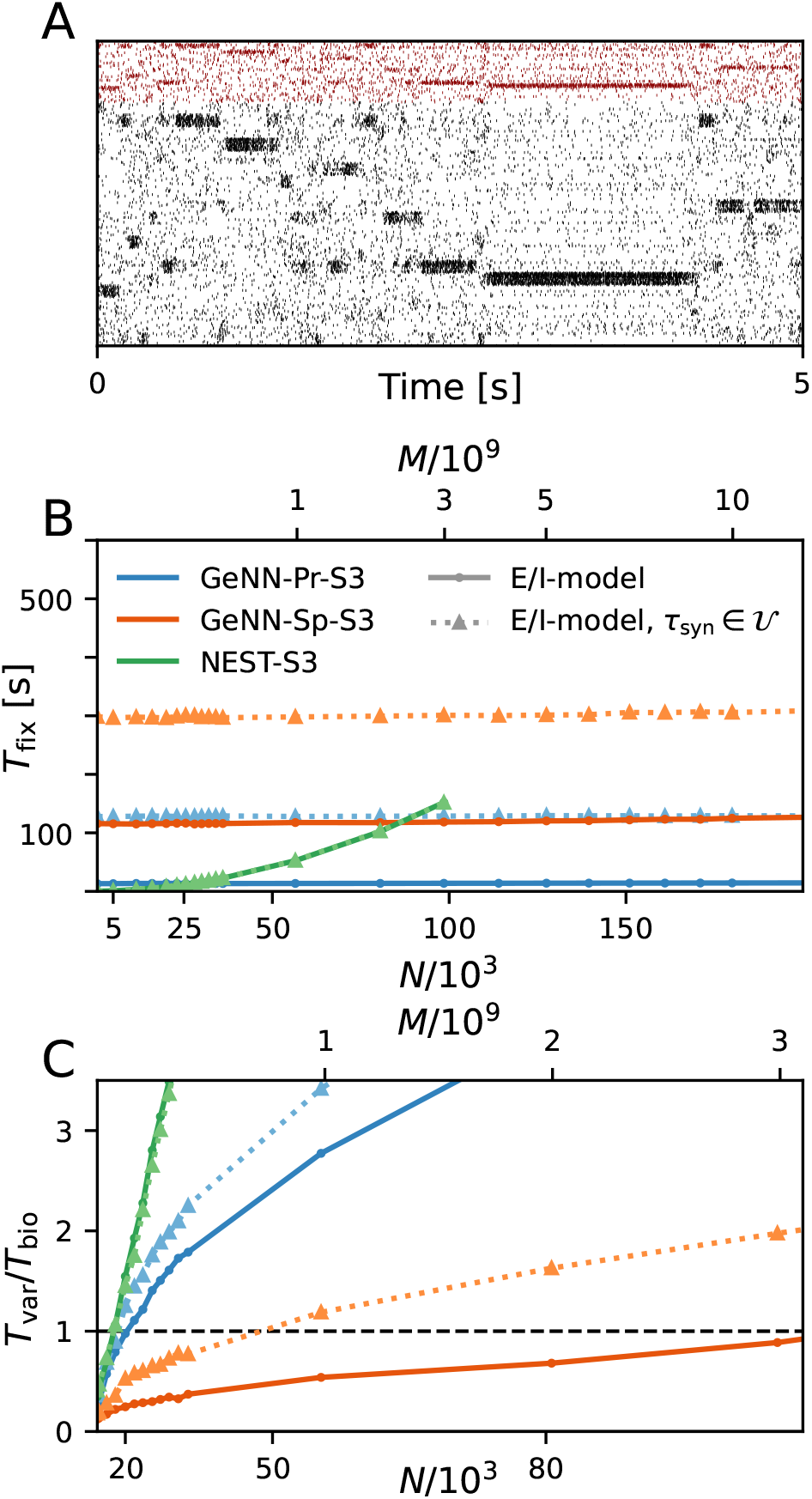
Metastability and simulation costs for networks with heterogeneous synaptic time constants. (A) Raster plot of excitatory (black) and inhibitory (red) spiking activity in a network of *N* = 25, 000 neurons with heterogeneous synaptic time constants during 5 s of spontaneous activity shows the desired metastable behavior where different individual clusters can spontaneously assume states of high and low activity. Shown is a portion of 8% of neurons from each of the 20 clusters. Network parameters are identical to Fig.1 D: *J*_E+_ = 2.75, *I*_thE_ = 1.6, *I*_thI_ = 0.9. The spike raster plot is generated from a GeNN simulation. (B) Total fixed costs in dependence of network size for the E/I model with (dotted lines) and without (solid lines) distributed synaptic time constants when simulated with NEST (green) or GeNN. For NEST the fixed costs are indistinguishable for both model variants. For GeNN the fixed costs are independent of network size but considerably higher with distributed synaptic time constants. (C) Real-time factor in dependence of network size. For NEST the variable costs are the same in the case of homogeneous and heterogeneous synaptic time constants while GeNN has increased variable costs in the heterogeneous case. Note that the x-axis in panel B is linear in *N*, while12the x-axis in panel C is linear in *M*.

For the variable costs, there is again no increase with NEST with the same linear dependence on network size as in the homogeneous case (Fig. 4C). This was to be expected as NEST, by default, stores one propagator for the excitatory and one for the inhibitory input per neuron. Thus, the per neuron integration of the postsynaptic current is performed identically to the homogeneous case. In GeNN, on the other hand, we use independent synapse models where each individual synapse has a different time constant. This requires to perform the integration over time independently. Hence, we observe a considerable increase in variable costs that follows the same convex dependency on network size as in the homogeneous case (Fig. 4C). For the duration of 10 s biological model time used here and for a network size of 50.000 neurons, the total costs with GeNN are higher than with NEST (Table 3, see Discussion).

**Table 3.** Fixed costs (*T*_fix_) and variable costs (*T*_var_) for simulating networks of size *N* = 50.000 during 10 s of biological model time. Top: standard benchmark model with clustered connectivity (*J*_*E*+_ = 10) and homogeneous synaptic time constants (E/I-model). Middle: random balanced network (RBN) without clustering (*J*_*E*+_ = 1) and homogeneous synaptic time constants. Bottom: clustered network model with (*J*_*E*+_ = 10) and with heterogeneous synaptic time constants (*τ*_syn_ ∈ *U*).

| **E/I-model** | $T_{\text{fix}}$ [s] | $T_{\text{var}}$ [s] |
| --- | --- | --- |
| GeNN-Pr-S3 | 13.6 | 24.9 |
| GeNN-Pr-S1 | 21.6 | 80.3 |
| GeNN-Sp-S3 | 117.3 | 5.2 |
| GeNN-Sp-S1 | 202.8 | 19.8 |
| NEST-S3 | 41.9 | 80.8 |
| NEST-S2 | 104.7 | 280.3 |
| **RBN** | $T_{\text{fix}}$ [s] | $T_{\text{var}}$ [s] |
| GeNN-Pr-S3 | 13.4 | 16.0 |
| GeNN-Sp-S3 | 116.6 | 3.3 |
| NEST-S3 | 41.7 | 26.4 |
| **E/I-model, $\tau_{\text{syn}} \in \mathcal{U}$** | $T_{\text{fix}}$ [s] | $T_{\text{var}}$ [s] |
| GeNN-Pr-S3 | 128.2 | 30.2 |
| GeNN-Sp-S3 | 298.9 | 10.3 |
| NEST-S3 | 41.9 | 78.5 |

### Efficient approach to parameter grid search

Achieving robust model performance requires the vital and computationally demanding step of model calibration with respect to independent model parameters (see Discussion). Generally, the total costs for a parameter optimization directly scale with the number of samples tested for the considered parameter combinations. In our spiking attractor benchmark model we have 22 independent parameters (cf. Tables 6 and 7). To this end, we perform a 2D grid search investigating the average firing rate across the entire population of excitatory neurons in dependence on two parameters: the constant background stimulation currents *I*_*x*E_ and *I*_*x*I_ measured in multiples of the rheobase current *I*_*x*X_ = *I*_thX_ · *I*_rheoX_ (Fig. 5). We sampled a grid of 40 × 40 parameter values and for each sample point our benchmark model is simulated for 10 seconds biological model time. This results in a total simulated biological model time of 16,000 seconds for a network size of *N* = 25, 000 neurons and *M* = 193 ·10^6^ connections. We obtain plausible spike raster plots for different combinations of the background stimulation of excitatory and inhibitory neuron populations and metastability emerges in a large parameter regime (Fig. 5A). The activity of the excitatory populations increases along the *I*_thE_-axis and decreases with stronger stimulation of the inhibitory neurons. The comparison of average firing rates across the excitatory population in simulations with NEST (Fig. 5B) and GeNN (Fig. 5C) shows only negligible differences due to the random network structure.

**Figure 5.**
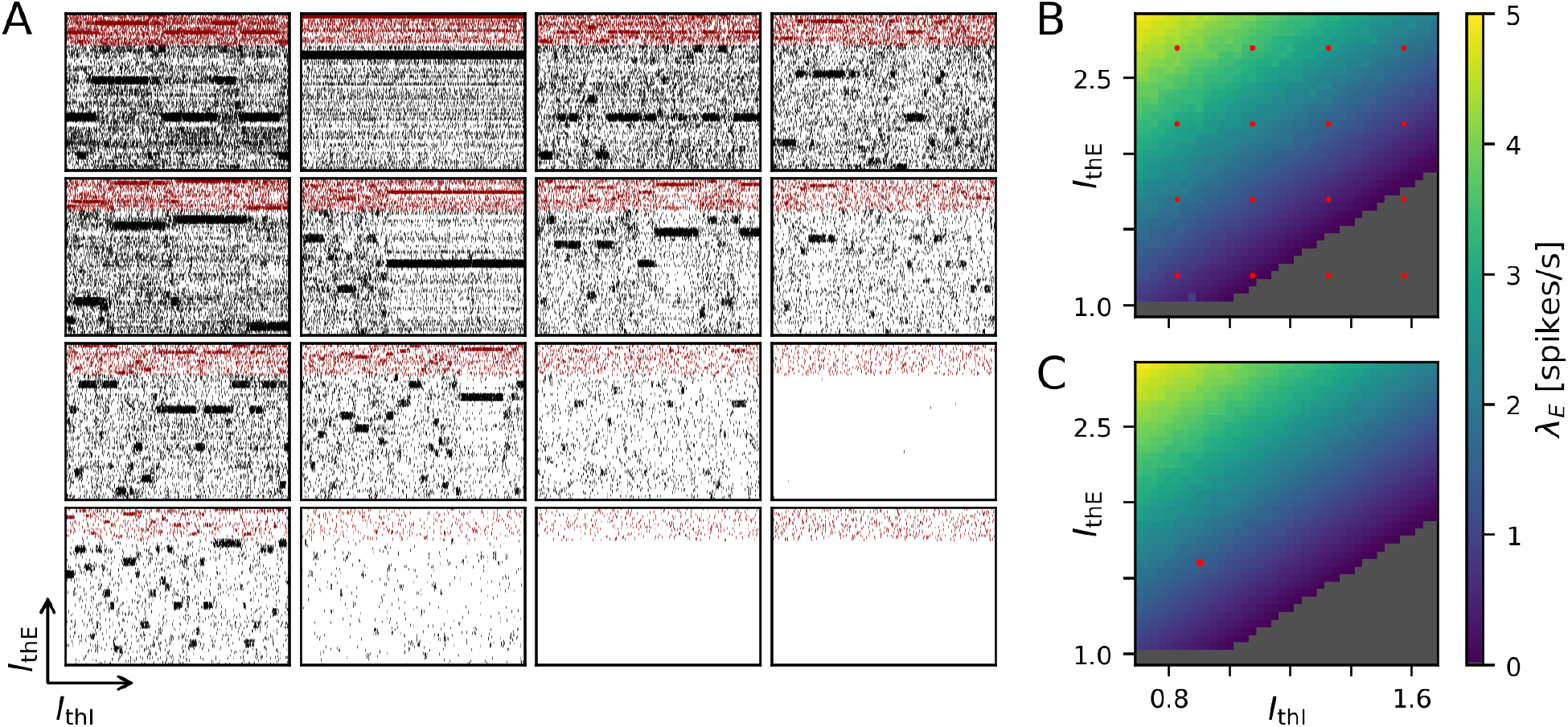
Two-dimensional parameter grid search with GeNN and NEST. (A) Shown are a subset of 4 × 4 spike raster plots as generated from simulations with NEST. The respective parameter values are indicated by the red point markers in B. In each raster 8% of neurons from each of the 20 clusters are displayed. The network size was *N* = 25, 000, and the clustering parameter was *J*_E+_ = 2.75 as in Fig. 1 D and Fig. 4. (B) Average spontaneous firing rates of all excitatory neurons in dependence of *I*_thE_ and *I*_thI_ across the complete grid of 40 × 40 parameter pairs when simulated with NEST. The run-time for completing the grid search was 29,692 seconds (≈ 8 h 15 min). (C) Same as in B for the simulation with GeNN in batch-mode (batch size 40). The red point marker indicates the parameter combination used for the spike raster plot in Fig. 1 D. The required run-time was 2,809 seconds (≈ 47 min).

We first compared this grid search for simulation with NEST on different servers and with different parallelization schemes (Table 4). For each parameter combination a new network instance was generated ensuring independent samples. The fastest grid search is achieved using server S3 (single CPU socket) with one worker that uses all available cores as threads. This parallelization scheme reduces the run-time by 40% in comparison to a scheme with six simulations run in parallel with four threads each. The observed advantage of using a single worker is in line with the results of (Kurth et al., 2022). The results are different on S2 with its two CPU sockets where 5 parallel simulations, each with 4 threads, resulted in a slightly improved performance compared to the case of a single simulation on all cores. Performing the same grid search with GeNN using independent network instances for each parameter combination required a total of 4.500 s and was thus 6.6 times faster than the independent grid search with NEST.

**Table 4.** Run-time of 40×40 grid search with GeNN and NEST with different hardware configurations and parallelization schemes. *n*_*w*_ denotes the number of workers and *n*_*T*_ the number of threads used for CPU simulation on multiple cores. S3* denotes the NEST simulation approach where a single network instance is set up once and reused for all samples in the grid.

| **GeNN** |  |  |
| --- | --- | --- |
| Server | Batch size | Run-time [s] |
| S3 | 40 | 2,809 |
| S3 | 1 | 4,500 |
| S1 | 4 | 13,080 |
| S1 | 1 | 14,491 |

| **NEST** |  |  |
| --- | --- | --- |
| Server | $n_W / n_T$ | Run-time [s] |
| S3* | 1 / 24 | 18,028 |
| S3 | 1 / 24 | 29,692 |
| S3 | 6 / 4 | 49,403 |
| S2 | 1 / 20 | 83,848 |
| S2 | 5 / 4 | 81,631 |

To maximize GPU utilization and to save fixed costs, we here propose an alternative batch mode for the parameter search with GeNN. It uses the same network connectivity for all instances in a batch and thus reduces memory consumption while using all cores of the GPU. To this end we distributed the instances of a single batch pseudorandomly across the entire grid. The identical connectivity introduces correlations across all network instances within a single batch (see Discussion) while a batch size of 1 results in fully independent networks. The shortest run-time was achieved for a batch size of 40 (Table 4) and resulted in a significant speed-up factor of 1.6 when compared to a batch size of one.

To match the batch mode in GeNN we tested an alternative approach with NEST in which we generate the network model only once and then re-initiate this same network for all parameter combinations in the grid. Now, network connectivity is identical across the complete parameter grid and thus not independent (see Discussion). In this approach, the reduced fixed costs for model setup (combined phases of node creation and connection) considerably reduced the overall wall-clock time for completing the grid search by almost 40 % (S3* in Table 4), resulting in a speed-up factor of 1.7 compared to the simulation of independent network instances. The batch-mode approach with GeNN was thus 6.4 times faster than the single network instance approach with NEST (S3*). We observed a considerable dependence of wall-clock time on the average firing rate for the grid search in NEST simulations. This dependence is comparably weak in the here tested `SPARSE` mode with GeNN.

## Discussion

### Limitations of the present study

We here restricted our benchmark simulations with NEST and GeNN to single machines with multiple CPU cores equipped with either a high-end or low-cost GPU (Table 1). We may consider this type of hardware configuration as standard equipment in computational labs. In our simulations with NEST on a single-processor machine (S3) we found that matching the number of threads to the number of cores was most efficient, in line with the results reported by Kurth et al. (2022), while on a dual-processor machine (S1) matching the number of threads and cores resulted in a small loss of speed compared to multiple simulations run in parallel (Table 4). We did not attempt to use the message passing interface (MPI) for distributing processes across the available cores. As pointed out by a previous study (Ippen et al., 2017), this can increase simulation performance with respect to simulation speed but at the same time considerably increases memory consumption, which would further limit the achievable network size.

NEST is optimized for distributed simulation by means of efficient spike communication across machines and processes (Pronold et al., 2022; Jordan et al., 2018; Ippen et al., 2017; Kunkel et al., 2014, 2012). This allows for scaling from single machines to multiple machines and allowed for the simulation of very large SNNs on supercomputers with thousands of compute nodes (Schmidt et al., 2018; Jordan et al., 2018; Kunkel et al., 2014). For the E/I cluster topology of our benchmark model, however, we do not expect a good scaling behavior of simulation speed in distributed environments for two reasons. One limiting factor for simulation speed is the communication of spikes between machines and the spike delivery on each machine. In NEST, communication between machines is optimized by communicating packages of sequential spikes, which requires a sufficiently large minimal synaptic delay (Morrison et al., 2005) and, thus, spike delivery on each machine dominates the cost of communication (Pronold et al., 2022; Jordan et al., 2018). Our current model implementation uses the minimum synaptic delay of only a single time step (0.1 ms). Second, the E/I cluster model shares the structural connectivity of the RBN (Fig. 1) where the topology of excitatory and inhibitory clusters is defined through connection strengths while connectivity is unaffected and comparably high with a pairwise connection probability of *p* ≈ 0.3. In future work we will consider an alternative structural definition of the cluster topology where the number of connections between E/I clusters is reduced, while it remains high within clusters. Simulating one or several E/I clusters on a single machine could then benefit distributed simulation due to a reduced spike communication between machines. A cluster topology defined by connectivity also opens the possibility to form clusters by means of structural plasticity (Gallinaro et al., 2022). We note that, in our current network definition and for large network size, the number of synapses per neuron exceeds biological realistic numbers in the order of 10,000 synapses per neuron were reported in the primate neocortex (Sherwood et al., 2020; Boucsein et al., 2011). However, here we deliberately used a fixed connectivity scheme across the complete investigated range of network sizes. The large number of synapses creates a high computational load for the spike propagation.

A number of GPU-based simulators are currently in use for SNN simulation, such as ANNarchy (Vitay et al., 2015), CARLsim (Niedermeier et al., 2022), BINDSnet (Hazan et al., 2018), GeNN and NEST GPU (Golosio et al., 2021). These use different design principles (Brette and Goodman, 2012; Vlag et al., 2019) that are optimal for specific use cases. NEST GPU, for example, follows the design principle of NEST allowing for the distributed simulation of very large networks on multiple GPUs (on multiple machines) using MPI. Simulation of the multi-area multi-layered cortical network model as defined in Schmidt et al. (2018) with a size of *N* = 4.13 · 10^6^ neurons has recently been benchmarked on different systems. Knight and Nowotny (2021) found that NEST simulation on the JURECA system (Thörnig, 2021) at the Jülich Supercomputing Centre was ≈ 15 times faster than simulation with GeNN on a single GPU (NVIDIA TITAN RTX). The recent work by Tiddia et al. (2022) found that NEST GPU (parallel simulation on 32 GPUs, NVIDIA V100 GPU with 16 GB HBM2e) outperformed NEST (simulated on JUSUF HPC cluster Von St. Vieth (2021) and JURECA) by at least a factor of two.

Kurth et al. (2022) reported the real-time factor for the multi-layered model of a single cortical column introduced by Potjans and Diesmann (2014) with about *N* = 80, 000 neurons and 0, 3 · 10^9^ synapses for a simulation with GeNN as *RT* = 0.7 (NVIDIA Titan RTX) and with NEST as *RT* = 0, 56 (cluster with 2 dual-processor machines with 128 cores each). For networks with approximately the same number of neurons and a higher number of connections we here report similar real-time factors. With GeNN-Sp-S3 we achieved *RT* = 0, 7 (cf. Fig. 3) for the E/I cluster network with *N* = 80, 500 neurons and 2 ·10^9^ synapses, and *RT* = 0, 56 (cf. Fig. S1) for the corresponding RBN. The faster simulation of the RBN results from a lower average firing rate of ≈ 0.9 spikes/s as compared to ≈ 8.5 spikes/s in the E/I cluster network.

Providing comparable benchmarks for the simulation of SNNs across different simulation environments and different hardware systems is generally hampered by two factors. First, different simulators use different design principles. To fully exploit their capabilities in a benchmark comparison one needs to optimize for each simulator and use case. Second, the community has not agreed on standardized benchmark models (Albers et al., 2022; Ostrau et al., 2022; Kulkarni et al., 2021; Steffen et al., 2021). The RBN is widely used in computational neuroscience. However, its definition varies considerably across studies e.g. with respect to connection probabilities, fixed versus distributed in-degrees of synaptic connections, neuron and synapse models, or the background stimulation by constant or noise current input. Second, SNN simulation environments are subject to continuous development affecting optimization for speed and memory consumption, which complicates comparability across different versions. We thus did not attempt to directly compare the run-time performance obtained for the model simulations considered in the present study to the performances reported in previous studies.

NEST provides a high degree of functionality, good documentation and many implemented neuron and synapse models. This results in a high degree of flexibility allowing, for instance, to introduce distributed parameters by using a NEST function that passes the respective distribution parameters as arguments when initializing the model, which is then set up from scratch. One obvious limitation of the flexibility of GeNN is the high fixed costs for model definition and building the model on the CPU and for loading the model on the GPU before it can be initialized. This limits its flexible use in cases where non-global parameters of a model change, which cannot be changed on the GPU. We would thus like to encourage the development of a method that automatically translates selected model parameters into GeNN variables for a given model definition, allowing to change those model parameters without recompilation between simulations. This functionality would allow to fully exploit the simulation speed on the GPU and benefit the time-to-solution while reducing the likelihood of implementation errors.

### Efficient long-duration and real-time simulation on the GPU

Our results show that GPU based simulation can support the efficient simulation over long biological model times (Fig. 3). This is desirable e.g. in spiking models that employ structural (Deger et al., 2012; Gallinaro et al., 2022) or synaptic (Vogels et al., 2011; Zenke and Ganguli, 2018; Sacramento et al., 2018; Asabuki et al., 2022; Illing et al., 2021) plasticity to support continual learning and the formation and recall of short-term (on the time scale of minutes), middle-term (hours) or long-term (days) associative memories. Similarly, simulating nervous system control of behaving agents in approaches to computational neuroethology may require biological model time scales of minutes to hours or days. The low variable simulation costs achieved with GeNN can also benefit real-time simulation of SNNs e.g. in robotic application.

We here considered spiking networks in the approximate size range from one thousand to a few million neurons. This range covers the complete nervous system of most invertebrate and of small vertebrate species as shown in Fig. 6 and Table 5, including for instance the adult fruit fly *Drosophila melanogaster* with *N*≈ 100, 000 neurons in the central brain (Raji and Potter, 2021), the European honeybee *Apis mellifera* (*N* ≈ 900, 000) (Witthöft, 1967; Godfrey et al., 2021; Menzel, 2012) and the zebrafish *Danio rerio* (≈ *N* 10 ·10^6^). In mammals, it covers a range of subsystems from a single cortical column with approximately 30, 000 −80, 000 neurons (Markram et al., 2015; Boucsein et al., 2011; Potjans and Diesmann, 2014) to the complete neocortex of the mouse (*N*≈ 5 · 10^6^ neurons, (Herculano-Houzel et al., 2013)). Models exceeding this scale are currently still an exception (Kunkel et al., 2014; Eliasmith and Trujillo, 2014; Van Albada et al., 2015; Jordan et al., 2018; Igarashi et al., 2019; Yamaura et al., 2020) and typically require the use of a supercomputer.

**Table 5.**
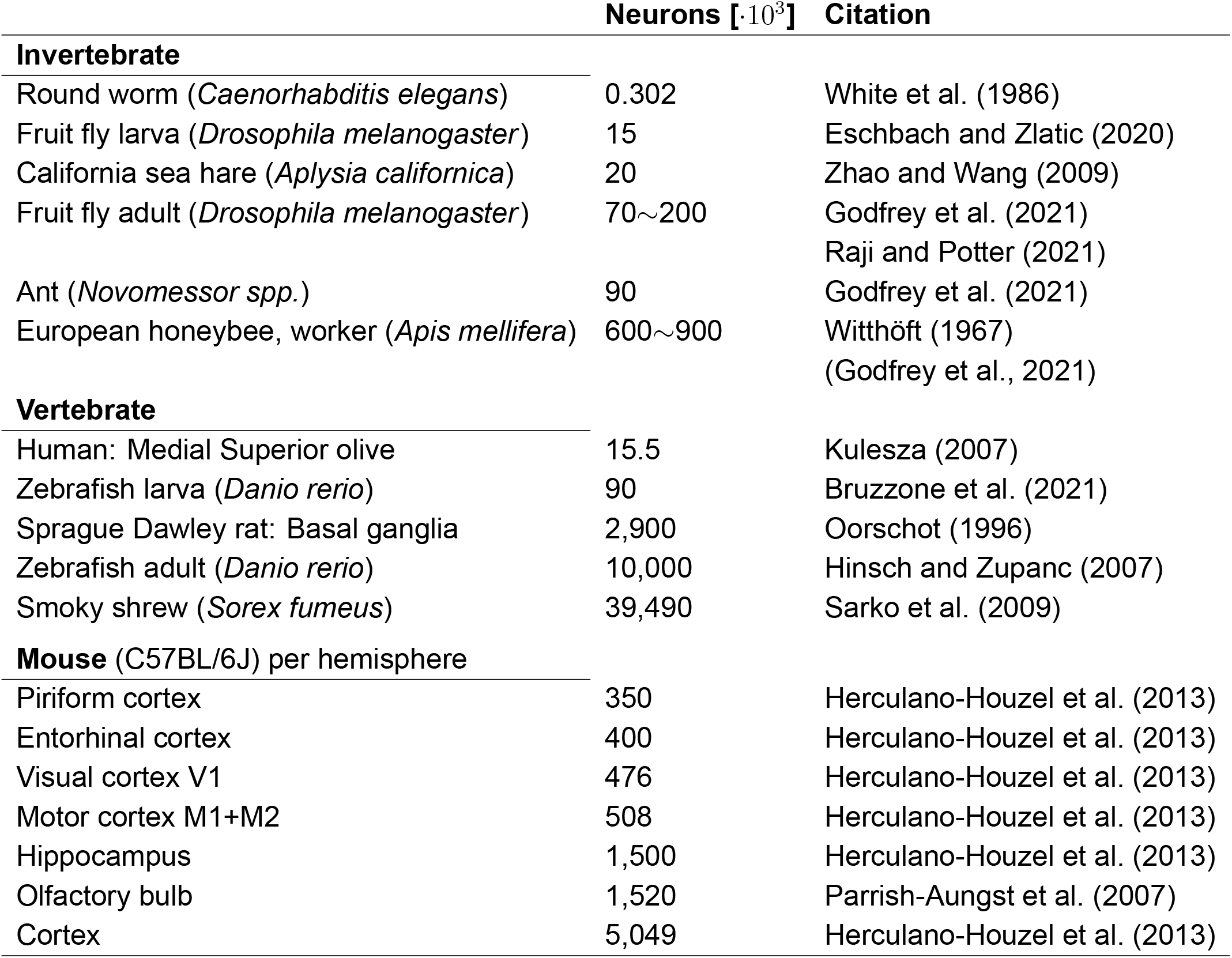
Number of neurons of selected model organisms and subsystems.

**Figure 6.**
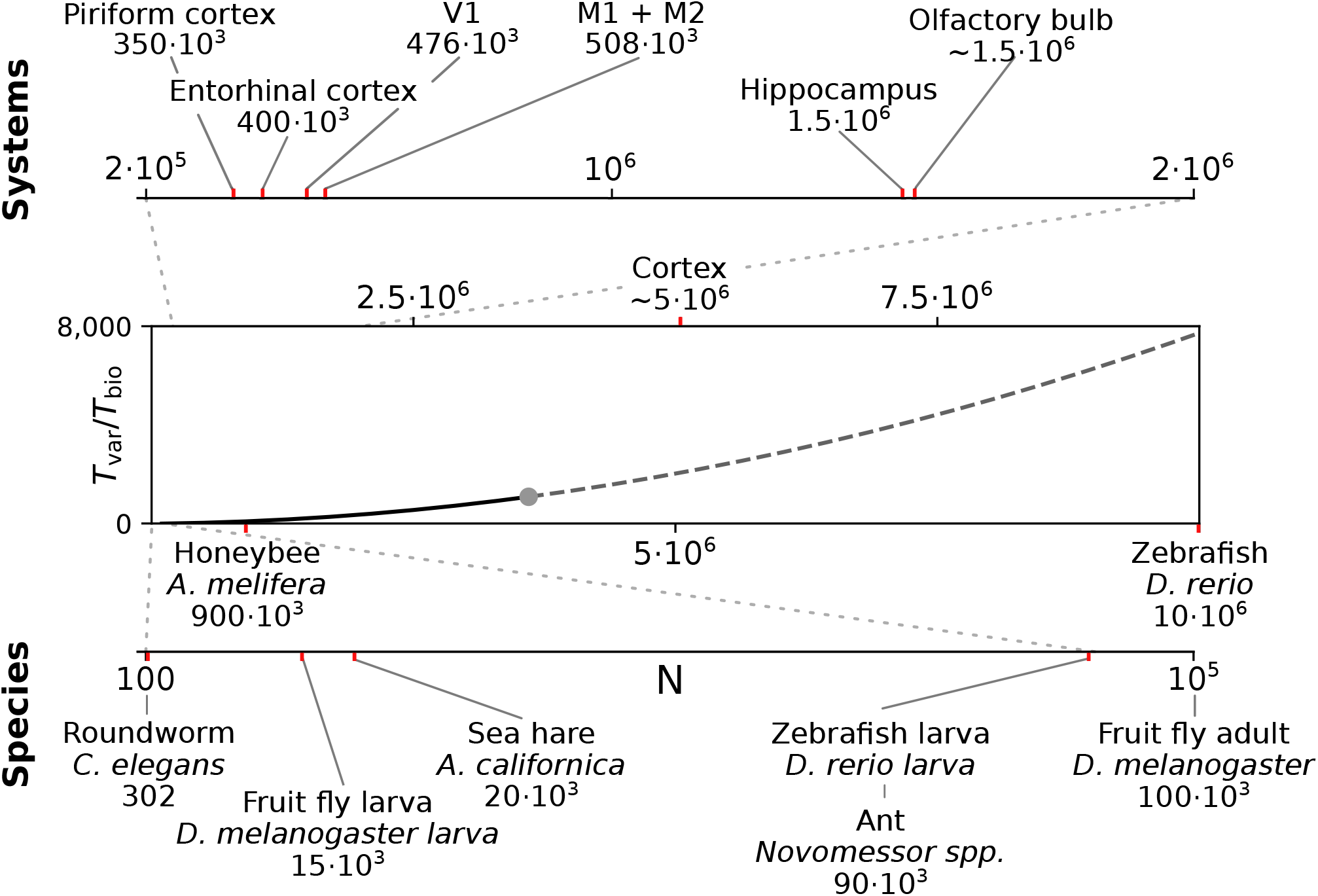
Full system modeling with spiking neurons. Wall-clock time relative to simulated biological model time *T*_*var*_ / *T*_*bio*_ over a realistic range of network sizes for spiking network simulation. Network size is projected to the total number of neurons per hemisphere for subsystems in the mouse (top), and in the CNS of invertebrate and small vertebrate model species (bottom); see also Table 5. The graph sketches the quadratic dependence on network size *N* for large *N* fitted for the PROCEDURAL connectivity simulation method with GeNN (Fig. 3, see Materials and Methods). The solid line shows the range of network sizes covered in our simulations (cf. Fig. 2) using a high-end GPU with 48 GB RAM (S3, Table 2). The dashed line extrapolates to a maximum size of 10 million neurons assuming larger GPU RAM that would allow to cover the neocortex of the mouse and the complete CNS of the zebrafish.

### Benchmarking with grid search

Spiking neural networks have a large set of parameters. These include connectivity parameters and parameters of the individual neuron and synapse models. Thus, both in scientific projects and for the development of real-world applications of SNNs, model tuning through parameter search typically creates the highest demand in computing time. Currently available methods (Feurer and Hutter, 2019) such as grid search or random search (LaValle et al., 2004; Bergstra and Bengio, 2012), Bayesian optimization (Parsa et al., 2019), and machine learning approaches (Carlson et al., 2014; Yegenoglu et al., 2022) require extensive sampling of the parameter space. We therefore suggest to include the parameter search in benchmarking approaches to the efficient simulation of large-scale SNNs.

To this end we exploited two features of GeNN: batch processing and the on-GPU initialization of the model. Batch processing was originally introduced to benefit the execution of machine learning tasks with SNNs. It enables the parallel computation of multiple model instances within a batch. In our example with a network size of *N* = 25, 000 and for simulating a biological model time of 10 s we obtained a speed-up factor of 1.6 for a batch size of 40 compared to a single model instance per run. The current version of GeNN requires that all model instances of a batch use identical model connectivity. Thus, quantitative results, e.g. of the average firing rates (Fig. 5), are correlated across all samples within one batch (for batch size > 1). We distributed the 40 instances of one batch pseudorandomly across the grid such that correlations are not systematically introduced among neighboring samples in the grid. An important future improvement of the batch processing that will allow for different connectivity matrices within a batch and thus for independent model connectivities is scheduled for the release of GeNN 5.0 (https://github.com/genn-team/genn/issues/463). The possibility to initialize and re-initialize a once defined model and connectivity on the GPU (Knight and Nowotny, 2018) uses the flexibility of the code generation framework. This allows to define, build and load the model to the GPU once and to repeatedly initialize the model on the GPU with a new connectivity matrix (per batch). It also allows for the variable initialization of e.g. the initial conditions of model variables such as the neurons’ membrane potentials. In addition, global parameters can be changed during runtime. After initialization of a model this allows, for example, to impose arbitrary network input as pre-defined in an experimental protocol.

We here propose that the batch processing with GeNN can be efficiently used not only to perform parameter search but also to perform batch simulations of the identical model with identical connectivity in parallel. This can be beneficial e.g. to generate multiple simulation trials for a given stimulation protocol allowing for across-trial statistics, or to efficiently generate responses of the same model to different stimulation protocols within a single batch. In NEST this can be efficiently achieved by re-initialization of the identical network (Table 4). We note that we have tested one additional alternative approach to the grid search with NEST where we set up the model connectivity once and, afterwards, for each initialization reconstructed the network from the stored connectivity. This lead to a significant reduction in performance for large networks (data not shown).

### Networks with heterogeneous neuron and synapse parameters

SNNs are typically simulated with homogeneous parameters across all neuron and synapse elements of a certain type. Using heterogeneous parameters that follow experimentally observed parameter distributions increases the biological realism of a model and has been argued to benefit model robustness and neuronal population coding (Gjorgjieva et al., 2016; Tripathy et al., 2013; Mejias and Longtin, 2012, 2014; Lengler et al., 2013; Litwin-Kumar et al., 2016). In the present study we performed benchmark simulations with heterogeneity in the single parameter of synaptic time constant to quantify its effect on simulation costs. The efficient solution provided by NEST for the specific neuron model used here does neither increase fixed costs nor variable costs (Fig. 4) in line with the results of Stimberg et al. (2019), albeit with the limitation to one single time constant per synapse type (excitatory and inhibitory) for each postsynaptic neuron. NEST offers an alternative neuron model that allows the definition of an arbitrary time constant for each synapse (see Materials and Methods) that was not tested in the present study. In GeNN, we had deliberately defined our neuron and synapse models separately (see Materials and Methods) as in future work we aim at introducing stochasticity of synaptic transmission to capture the inevitable variability of synaptic transmission in biology (Nawrot et al., 2009; Boucsein et al., 2011) that has been argued to support efficient population coding (Lengler et al., 2013).

### Metastability emerges robustly in attractor networks with large E/I clusters and heterogeneous synaptic time constants

With respect to attractor network computation, an important question is whether the functionally desired metastability can be reliably achieved in large networks and for large population sizes of neuron clusters. In our previous work we had limited our study of attractor networks to a maximum network size of 5, 000 neurons and could show that the topology of excitatory-inhibitory clusters benefit metastability for a varying number and size of clusters while pure excitatory clustering failed to support metastability for larger cluster size (Rostami et al., 2022; Rost et al., 2018). In our calibration approach of Fig. 5 we simulated networks of *N* = 25, 000 with a cluster size of 1, 000 excitatory and 250 inhibitory neurons. This results in the robust emergence of metastable activity for a reasonable regime of excitatory and inhibitory background currents. Metastability was retained when introducing distributed excitatory and inhibitory time constants (Fig. 4). We hypothesize that, due to its local excitation-inhibition balance (Rostami et al., 2022), the E/I cluster topology affords metastability for very large network and cluster sizes.

## Materials and Methods

### Hardware configurations

We perform benchmark simulations on hardware systems that can be considered as standard equipment in a computational research lab. We did not attempt to use High Performance Computing facilities that, for most users, are available only for highly limited computing time and require an overhead in scheduling simulation jobs. We employ three compute server systems specified in Table 1, which were acquired between 2016 and 2022. The amount of investment at the time of purchase has been fairly stable on the order of $7, 000− $10, 000 depending on whether a state-of-the-art GPU was included. Servers S1 and S3 are equipped with GPUs. The GeForce GTX 970 (S1; NVIDIA, Santa Clara, USA) can now be considered a low-cost GPU in the price range of $ 300. The Quadro RTX A6000 (S3; NVIDIA, Santa Clara, USA) is one example of current state-of-the-art high-end GPUs, for which prices vary in the range of $3, 000 −$5, 000. We use the job scheduler HTCondor (Thain et al., 2005) on all servers independently with one job scheduler per server. We ensure acquisition of all cores and the complete GPU to a running job and prevent other jobs from execution till the running job is finished.

### Simulators

We benchmark with the Neural Simulation Tool (NEST) (Gewaltig and Diesmann, 2007) and the GPU-enhanced Neuronal Networks (GeNN) framework (Yavuz et al., 2016) that follow different design principles and target different hardware platforms. The Neural Simulation Tool (NEST) (Gewaltig and Diesmann, 2007) is targeted toward computational neuroscience research and provides various biologically realistic neuron and synapse models. Since the introduction of NESTML (Plotnikov et al., 2016) it also allows custom model definitions for non-expert programmers. NEST uses the so-called simulation language interpreter (SLI) to orchestrate the simulation during run-time as a stack machine. This allows a modular design of the whole simulator and thus the usage of pre-compiled models. NEST supports parallelization across multiple threads via Open Multi-Processing (OpenMP) as well as multiple processes, which can be distributed across multiple machines via the message passing interface (MPI). It is suitable for the whole range of desktop workstations to multi-node High-Performance clusters. We use NEST version 3.1 (Deepu et al., 2021) (in its standard cmake setting) with the Python interface PyNEST (Eppler et al., 2009) and Python version 3.8 to define our model and control the simulation.

GeNN is a C++ library to generate code from a high level description of the model and simulation procedure. It employs methods to map the description of the neuron models, the network topology as well as the design of the experiment to plain C++ code, which then is build for a specific target. GeNN supports single-threaded CPUs or a GPU with a CPU as host. The scope of GeNN is broader than that of NEST. With features like the batch-mode, which allows for inference of multiple inputs, GeNN becomes especially useful for machine learning tasks with SNNs as well as for general research in computational neuroscience. GeNN is more rigid in the network topology and in its parameters after the code generation is finished. It does not support general reconfiguration of the network during the simulation. If GeNN is used on a GPU, the CPU is used to generate and build the simulation code and orchestrate the simulation. All other time consuming processes such as initialization of the connectivity and variables of the model, update of state variables during simulation, and spike propagation are performed by the GPU. The model construction in the GPU memory is run by loading the model or can be rerun by reinitializing the model, which affects the connectivity and state variables as well as the spike buffers. The code generation framework in GeNN allows for a heavily optimized code depending on the use case. One of these optimization possibilities is the choice of connectivity matrices. Here, we utilize only the `SPARSE` and the `PROCEDURAL` connectivity as explained below.

#### SPARSE connectivity

The connectivity matrix is generated on the GPU during the loading of the model and if the model is reinitialized. It is persistent during the simulation. No additional computational load is generated during the simulation.

#### PROCEDURAL connectivity

Single elements of the connectivity matrix are generated on demand during simulation for the spike propagation. Only fixed random seeds are saved during the loading of the model and if the model is reinitialized. An additional computational load is caused during the simulation, but the memory consumption is low.

Furthermore we use global synapse models for all simulations except for simulations with distributed synapse parameters. In simulations with distributed synaptic time constants we use synapse models with individual postsynaptic model variables. We use the released GeNN version 4.6.0 and a development branch of version 4.7.0, which is now merged into the main and released (commit ba197f24f), with its Python interface PyGeNN (Knight et al., 2021) and Python version 3.6.

### Neuron models and network architectures

We use the spiking neural network described by Rostami et al. (2022) as benchmarking model. This model uses leaky integrate-and-fire neurons with exponentially shaped postsynaptic currents. The subthreshold dynamics evolves according to

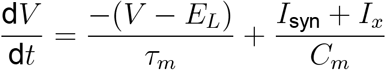

and the synaptic current to a neuron i evolves according to

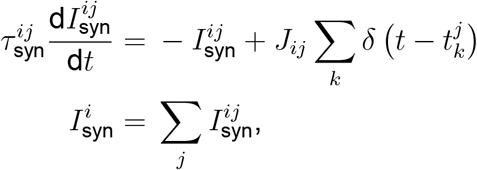

where 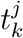 is the arrival of the *k*^th^ spike of the presynaptic neuron *j* and *δ* is the Dirac delta function.

We use the NEST model `iaf_psc_exp`, which employs an exact integration scheme as introduced by Rotter and Diesmann (1999) to solve the above equations efficiently for two different synaptic input ports. The input ports can have different time constants. We implement the same integration scheme in GeNN but modify it to fit only one synaptic input port while treated as piece-wise linear to combine different synapse types in terms of their time constant.

We follow the widely-used assumption that 80% of neurons in the neocortex are excitatory while 20% are inhibitory to build the network model, which is based on the statistical work in the mouse neocortex by Braitenberg and Schüz (1998). Connections between neurons are established with the probabilities *p*_EE_ = 0.2, *p*_EI_ = *p*_IE_ = *p*_II_ = 0.5. Autapses and multapses are excluded. Excitatory and inhibitory populations are divided into *N*_*Q*_ clusters by increasing the weights of intra-cluster connections while decreasing the weights of intercluster connections. Weights are calculated to ensure a local balance between excitation and inhibition, as introduced before for binary networks (Rost et al., 2018). Parameters are given in Tables 6 and 7. For comparability we matched our parameters to those used in previous related works (Litwin-Kumar and Doiron, 2012; Mazzucato et al., 2015; Rost, 2016; Rostami et al., 2022). Synaptic weights *J*_XY_ as well as the background stimulation currents are scaled by the membrane capacitance *C*. By this scaling, the capacitance has no influence on the dynamics of the network but only on the magnitude of PSCs and the background stimulation currents. Following previous studies, we set the value of capacitance to *C*= 1pF.

**Table 6.**
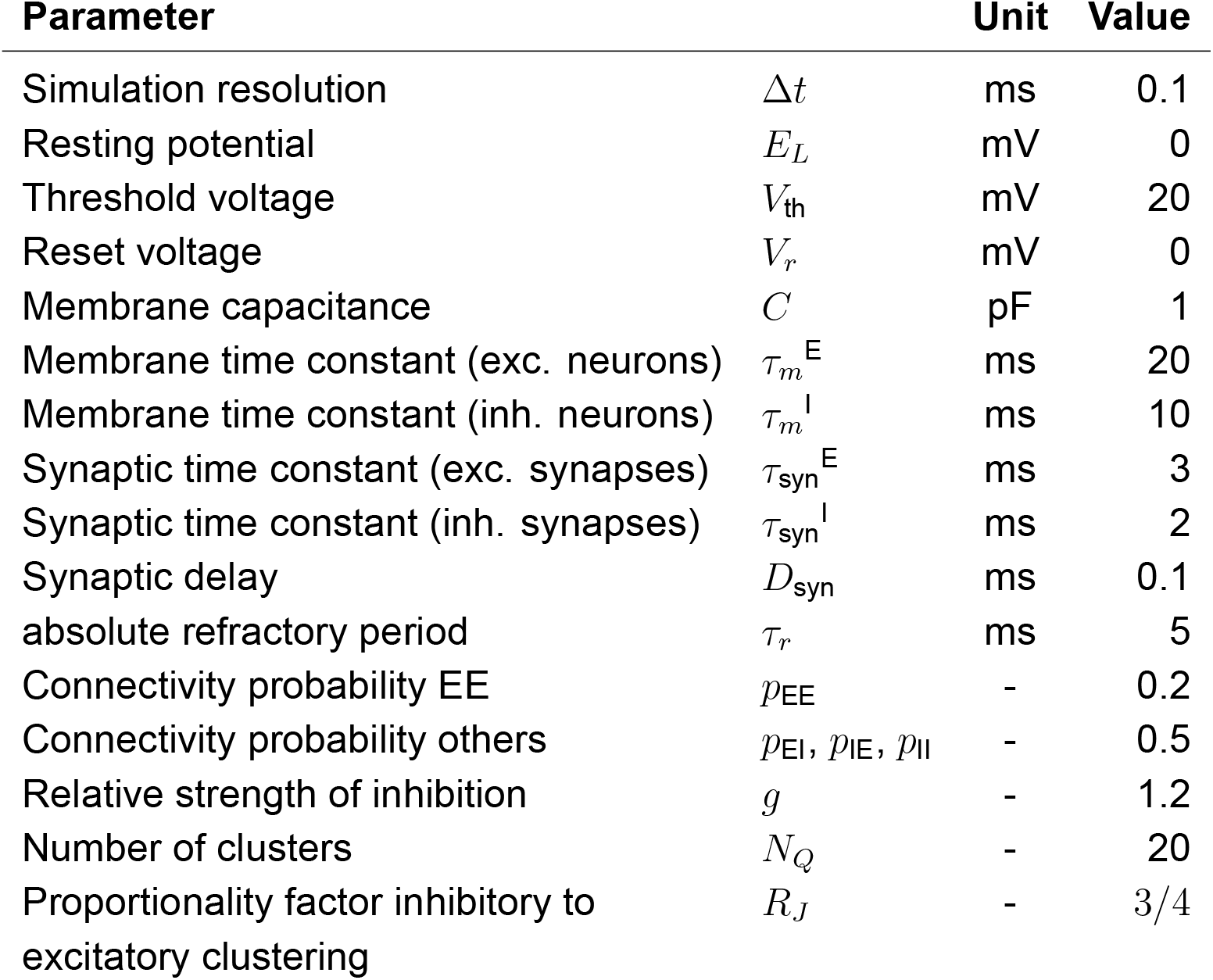
Constant model parameters for all simulation modalities.

| Parameter |  | Unit | Value |
| --- | --- | --- | --- |
| Simulation resolution | $\Delta t$ | ms | 0.1 |
| Resting potential | $E_L$ | mV | 0 |
| Threshold voltage | $V_{th}$ | mV | 20 |
| Reset voltage | $V_r$ | mV | 0 |
| Membrane capacitance | $C$ | pF | 1 |
| Membrane time constant (exc. neurons) | $\tau_m^E$ | ms | 20 |
| Membrane time constant (inh. neurons) | $\tau_m^I$ | ms | 10 |
| Synaptic time constant (exc. synapses) | $\tau_{syn}^E$ | ms | 3 |
| Synaptic time constant (inh. synapses) | $\tau_{syn}^I$ | ms | 2 |
| Synaptic delay | $D_{syn}$ | ms | 0.1 |
| absolute refractory period | $\tau_r$ | ms | 5 |
| Connectivity probability EE | $p_{EE}$ | - | 0.2 |
| Connectivity probability others | $p_{EI}, p_{IE}, p_{II}$ | - | 0.5 |
| Relative strength of inhibition | $g$ | - | 1.2 |
| Number of clusters | $N_Q$ | - | 20 |
| Proportionality factor inhibitory to excitatory clustering | $R_J$ | - | 3/4 |

**Table 7.** Variable model parameters for different simulation modalities. *f* (*N*) denotes the dependency of the synaptic weights (*J*_XY_) on the network size. *J*_E+_ is the clustering strength of the excitatory neurons and influences the inhibitory clustering strength via the proportionality factor *R*_*J*_. Implementation details are described in Rostami et al. (2022).

| Parameter |  |  | Value |  |  |  |
| --- | --- | --- | --- | --- | --- | --- |
|  |  |  | 5,000 | 50,000 | Var N | Grid |
| Number exc. neurons | $N_E$ | - | 4,000 | 40,000 | $0.8 \cdot N$ | 20,000 |
| Number inh. neurons | $N_I$ | - | 1,000 | 10,000 | $0.2 \cdot N$ | 5,000 |
| Current stimulation exc. neurons | $I_{xE}$ | pA | 2.13 | 2.13 | 2.13 | 0.95 - 2.9 |
| Current stimulation inh. neurons | $I_{xI}$ | pA | 1.24 | 1.24 | 1.24 | 0.7 - 1.675 |
| Clustering strength exc. neurons | $J_{E+}$ | - | 10.0 | 10.0 | 10.0 | 2.75 |
| Synaptic weight EE (RBN) | $J_{EE}$ | pA | 0.329 | 0.104 | $f(N)$ | 0.147 |
| Synaptic weight IE (RBN) | $J_{IE}$ | pA | 0.250 | 0.079 | $f(N)$ | 0.112 |
| Synaptic weight EI (RBN) | $J_{EI}$ | pA | -0.877 | -0.277 | $f(N)$ | -0.392 |
| Synaptic weight II (RBN) | $J_{II}$ | pA | -1.337 | -0.423 | $f(N)$ | -0.598 |

In our model, each neuron can be presynaptic and postsynaptic partner to all other neurons. As a result, the number of synapses *M* scales quadratically with the neuron number *N*. We calculate the expectation of the number of synapses *M* by using the assumption of the portioning into 80% excitatory and 20% inhibitory neurons and calculating the expected number of synapses between the combinations of both by using the connection probabilities. Due to our exclusion of autapses we have to reduce the number of postsynaptic possibilities by one for the connections among excitatory neurons as well as among inhibitory neurons. The overall network connectivity *p* = 0.308 is determined by

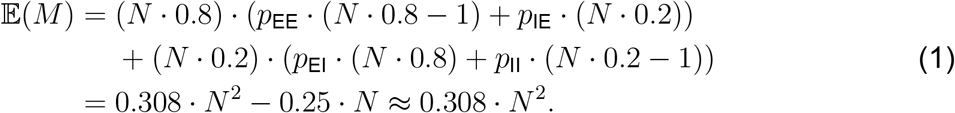

Additionally, we implement a model with excitatory and inhibitory synaptic time constants drawn from a uniform distribution with the same means as provided in Table 6 and a standard deviation of 5% of the mean. In NEST this is achieved by using the `nest.random.uniform` function as argument for the parameters `tau_syn_ex` and `tau_syn_in` of the neuron model. This results in distributed synaptic time constants across neurons. Thus, all synaptic inputs of one input type (excitatory and inhibitory) to a postsynaptic neuron have the same time constant. Replacing the neuron model by a multisynapse neuron model (e.g. `iaf_psc_exp_multisynapse`) would enable to use different time constants for subsets of the synaptic inputs of a single neuron. In GeNN we implement the distribution of synaptic time constants by re-implementing the postsynaptic model of an exponential PSC. We provide the decay factor exp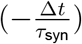 as variable to minimize the calculations during the simulation. GeNN only allows random number initialization for variables and not for parameters. Parameters within a group (neurons as well as synapses are generated as groups) have to be the same in GeNN. To initialize the decay factors corresponding to the uniform distribution of synaptic time constant, we define a custom variable initialization method. The generated network model does not enforce the same synaptic time constants for all inputs to a neuron. Deviating from the NEST model, the synaptic time constants are thus distributed across all connections rather than across neurons. A limited distribution across neurons as in NEST could be implemented in GeNN by implementing the synapse dynamics in the neuron model.

### Simulations

We perform two different sets of simulations with all tested combinations of a server and a simulator. We enforce a maximum run-time of five hours per simulation. The first set contains simulations of two networks with the sizes of 5,000 and 50,000 neurons and simulation times of 0; 1; 5; 25; 50; 100; 175; 350; 625; 1,250 and 2,500 seconds biological model time (zero seconds is used to determine the fixed costs for simulation preparation). We discard 10% as presimulation time from our analyses of network activity. We execute each simulation 5 times with different seeds of the random number generator. All simulations in the second set are 10 seconds long (biological model time; 1 second presimulation time and 9 seconds simulation time) and we used network sizes between 500 neurons and ≈ 3, 6 million neurons. We execute each simulation 10 times with different seeds of the random number generator. Based on the recorded wall-clock times, we determine the maximum network size for each configuration that fulfills real-time requirements.

For GeNN we use the spike recorder, which saves the spikes during the simulation in the RAM of the GPU and allows a block transfer of all spikes for a given number of simulation steps. The consumption of GPU-RAM is composed of the model with all its state variables and connectivity matrix, and of the memory for the spike recorder. The memory consumption of the model is independent of the simulated biological model time, but heavily dependent on the network size and the choice of the connectivity matrix type. The size of the memory of the spike recorder depends on the network size and the number of simulation steps, and thus on the simulated biological model time. We implement a partitioning of simulations into sub-simulations, if the GPU-RAM is too small to fit the model and the spike recorder in one simulation. We determine the maximum number of simulation steps for a large network that fits in the GPU-RAM and use this number as a heuristic. If the product of network size and simulation steps exceeds the heuristic, we iteratively divide the simulation into two until the product of both is smaller than the heuristic. As a fallback mechanism we include an error handling to divide the simulation further, if the model and the spike recorder do not fit the GPU-RAM. We transfer the spikes from the GPU to the host after each sub-simulation and process them into one list containing the neuron-ID and the spike time by a thread pool with a size of 16 workers. A greater number of workers exceeded the available RAM on our servers for large networks. All simulations are performed once with the `SPARSE` connectivity format and once with the `PROCEDURAL` connectivity. We delete the folder with the generated and compiled code before we submit the first job to the scheduler in order to prevent the use of code from previous code generation runs. We run one complete sequence of all different biological model times before switching to the other matrix type. This ensures that the code is generated five and ten times respectively for the two different simulation sets.

We use the same order of execution for our simulations in NEST. We match the number of OpenMP threads of the simulations in NEST to the number of cores available *n*_*T*_ = *n*_cores_ on the server. We do not utilize parallelization with MPI to minimize memory consumption (Ippen et al., 2017) and despite possible advantages in speed.

We additionally simulate the E/I cluster network model with distributed synaptic time constants with both simulators on S3 as described above. The network with *J*_E+_ = 1 represents the random balanced network (RBN) with constant background current stimulation. For this we use the same code and parameters as for the clustered network model for all calibrations.

### Grid search

We implement a general framework to perform a grid search. The framework takes multiple parameters and generates a regular mesh grid of the parameter values. It simulates each point in the grid, analyzes the network activity in the simulation, and saves the result together with its position in the grid to a pickle file. Pickle is a module from the Python Standard Library, which can serialize and de-serialize Python objects and save them to binary files. Pickle allows saving multiple objects in one file. Simulations are performed with the same network definition as used for the simulations before. All simulations of individual samples are 10 seconds long (biological model time; 1 second presimulation time and 9 seconds simulation time). We simulate a 2D grid with 40 × 40 samples.

We implement the grid search in GeNN by utilizing the batch-mode with *n*_Batch_ network instances in each run. We add global parameters to the model description. Global parameters are not translated into constants during compilation, but can be set independently for each instance in a batch and can be modified during run-time. The instances share all other parameters and the specific connectivity matrices. We use the `SPARSE` connectivity format and generate the simulation code only one time and then reinitialize the network after each batch and set the global parameters to their respective values. The reinitialization regenerates the connectivity matrix and resets all state variables as well as the spike buffer. The parameters as well as the batch size *n*_batch_ can only be changed by recompilation. Each time a number of samples equal to *n*_batch_ is drawn without replacement from the grid until all points in the grid are simulated. If fewer samples as *n*_batch_ are left, the free slots are filled by setting the global parameters to 0; the results of these slots are ignored. After a simulation the spike times are transferred in a block transfer and are processed by *n*_*W*_ = 24 workers into a matrix with the size 2 × *n*_spikes_ × *n*_batch_, where the first row contains the spike times, the second row the ID of the neuron, which emitted the spike and the third dimension corresponds to the different network instances in the batch. We use the same representation of the spike times and the senders for GeNN and NEST. The processing takes a reasonable amount of computation time as GeNN implements the spike recorders in the neuron populations and returns no global IDs but instead IDs for the neurons in the specific population. Afterwards, we analyze the spike times serially for each instance with the specific analyses, which are the firing rates of the excitatory and inhibitory populations in this grid search. The process writes the results afterward to the pickle file. We test the grid search with different batch sizes on S1 and S3 (see Table 4).

We implement the grid search in NEST by creating a list of all parameters in the grid and then parallelizing the simulations with Pathos by *n*_*W*_ workers. The workers import NEST independently, thus each simulation is completely independent. Each worker utilizes *n*_*T*_ threads. We match the number of cores available on the system: *n*_cores_ = *n*_*W*_ · *n*_*T*_. The workers write the results to one pickle file, which is protected by a lock to ensure data integrity by only allowing one worker to write at a time. This pickle file does not contain by default the spike times but instead only the result of the specific analyzes applied. We test different combinations of *n*_*W*_ and *n*_*T*_ on S2 and S3 (see Table 4).

We extend the NEST implementation for grid search by allowing the re-use of a once set up network. Before each run, the spike recorder is reset to zero events, the membrane voltages are reinitialized and the parameters are set for the current run. All runs thus use the same network with fixed connectivity.

### Definition of fixed and variable simulation costs

We divide the simulation cost into fixed and variable costs:

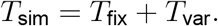

The fixed costs involve all steps necessary to set up and prepare the model before the first simulation step. They are independent of the biological model time to be simulated. The variable costs involve the actual propagation of the network state during each time step and the overall wall-clock time used for data collection during a simulation. We include timestamps in the simulation scripts to access the run-time of different execution phases. Due to the different design principles, not all phases of GeNN can be mapped to NEST. Table 8 shows the defined phases and the commands, which are executed in each phase. We included in the model definition phase of GeNN all steps necessary to set up the model description and the simulation itself. This contains the definition of neurons, their connectivity, the setup of the spike recorders, the stimulation of neurons and the definition of the neuron parameters as well as the parameters of the simulation itself such as the time resolution. During the build phase in GeNN, the code generation is executed on the basis of the defined model and the code is compiled for its target, which in our case is the specific server with a GPU. This process is not necessary for NEST because of its design using an interpreter to orchestrate the simulation by using pre-compiled models. The load phase in GeNN contains the construction of the model and all needed procedures to simulate it in the host memory and in the GPU-memory, and the initialization of the model. This includes the connectivity, variables of the model and variables of the general simulation. The initialization of the model can be also re-run manually to generate a fresh model based on the currently loaded code, as used for the grid search with GeNN. The model setup in NEST comprises the node creation and the connection phase. The former creates the structures of neurons, spike recorders and stimulation devices, and the general simulation parameters like the the parallelization scheme. In the latter the structures of the connectivity of nodes is created. Due to the ability of NEST to use distributed environments, presynaptic connectivity structures can only be created after the calibration of the system. This calibration contains the determination of delays between MPI processes if used, the allocation of buffers and calibrating nodes. The simulation phase contains for both simulators the propagation of the state and the delivery of spikes. In the case of GeNN each call of the simulate function only simulates one time step and thus the call has to be issued as many times as needed to complete the simulation. In the case of NEST this is done automatically. The download phase contains the query of the recorded spikes and the generation of our used format of representation as described for the grid search. In the case of GeNN, the conversion is proceeded by the transfer of the recorded spikes from the GPU to the host.

**Table 8.**
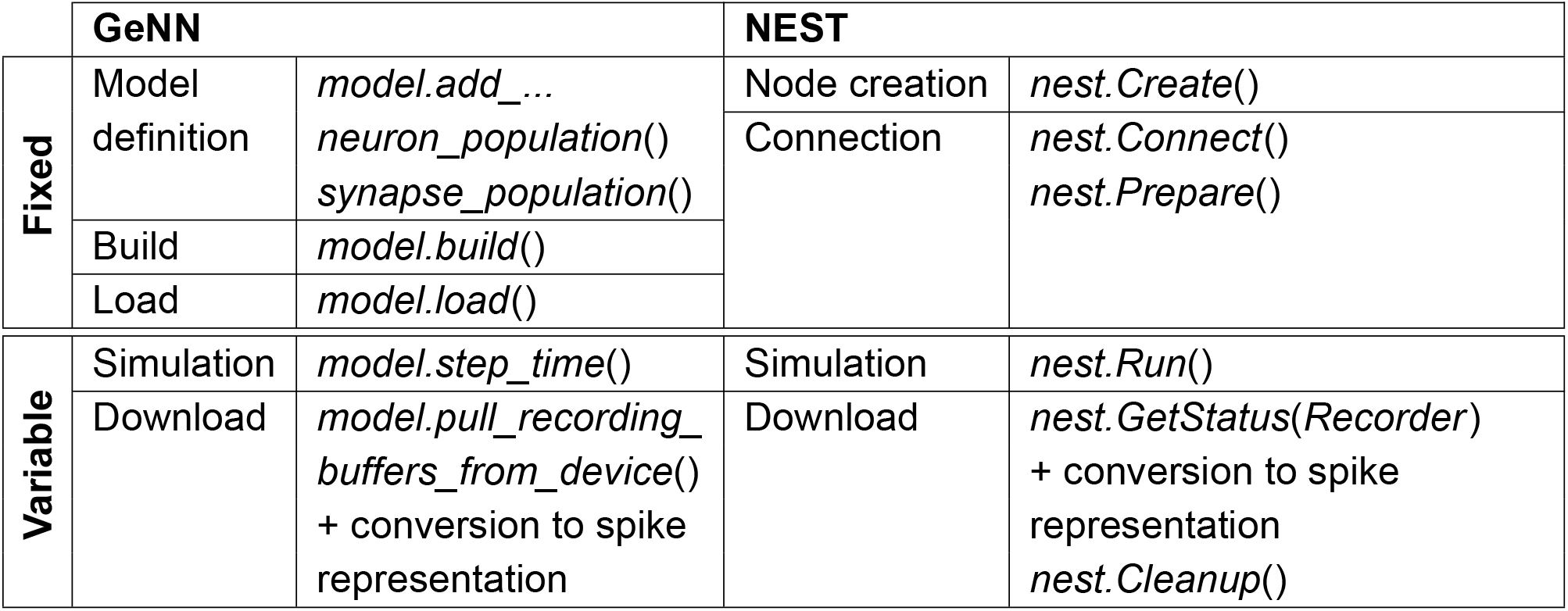
Execution phases of simulation

We report the median values of fixed and variable simulation costs across repeated simulations.

### Extrapolation to large network sizes

We can approximate the parabolic dependence of the proportionality factor RT = *T*_var_/*T*_bio_ on the number of neurons *N* for the `PROCEDURAL` simulation approach with GeNN (Fig. 3B and C) by the polynomial function

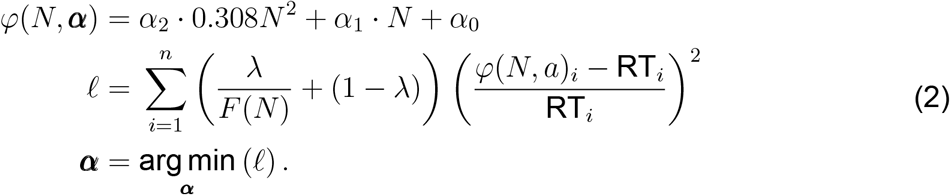

To this end we used a weighted least squares fit of Eq. 2 with three modifications: *i*) We scale the quadratic term by the connection density *p* = 0.308 to relate this term to the number of synapses *M, ii*) we use the relative error, and *iii*) we weight the samples by the inverse of the density along the independent variable *N*. This balances the influence of the large network with a smaller number of samples and the larger number of samples for small networks with the factor *λ* = 0.75. We estimate the density *F* (*N*) by a kernel density estimation using a Gaussian kernel (*σ* = 10, 000 neurons). The resulting fit is used for the extrapolation of *T*_var_/*T*_bio_ to larger network sizes as shown in Fig. 6. Simulation of larger networks will require larger GPU RAM.

## Author Contributions

FS and MN designed the research. FS carried out simulations and analysis of results. VR and MN supervised project. FS and MN wrote the manuscript.

## Acknowledgments

We are grateful to James Knight for his support with GeNN.

## Data Availability Statement

The generated data and the source code to produce this data as well as scripts to analyze it can be found in the GitHub repository https://github.com/nawrotlab/SNN_GeNN_Nest.

## References

Abeles, M. (1991). Corticonics: Neural circuits of the cerebral cortex. Cambridge University Press.

Albers, J., Pronold, J., Kurth, A. C., Vennemo, S. B., Haghighi Mood, K., Patronis, A., Terhorst, D., Jordan, J., Kunkel, S., Tetzlaff, T., Diesmann, M., and Senk, J. (2022). A Modular Workflow for Performance Benchmarking of Neuronal Network Simulations. Frontiers in Neuroinformatics, 16:837549.

Amit, D. J. and Brunel, N. (1997). Model of global spontaneous activity and local structured activity during delay periods in the cerebral cortex. Cerebral Cortex, 7(3):237–252.

Asabuki, T., Kokate, P., and Fukai, T. (2022). Neural circuit mechanisms of hierarchical sequence learning tested on large-scale recording data. PLOS Computational Biology, 18(6):1–25.

Bartolozzi, C., Indiveri, G., and Donati, E. (2022). Embodied neuromorphic intelligence. Nature Communications, 13:1024.

Bekolay, T., Bergstra, J., Hunsberger, E., DeWolf, T., Stewart, T., Rasmussen, D., Choo, X., Voelker, A., and Eliasmith, C. (2014). Nengo: a Python tool for building large-scale functional brain models. Frontiers in Neuroinformatics, 7.

Ben-Shalom, R., Ladd, A., Artherya, N. S., Cross, C., Kim, K. G., Sanghevi, H., Korngreen, A., Bouchard, K. E., and Bender, K. J. (2022). NeuroGPU: Accelerating multi-compartment, biophysically detailed neuron simulations on GPUs. Journal of Neuroscience Methods, 366:109400.

Bergstra, J. and Bengio, Y. (2012). Random search for hyper-parameter optimization. The Journal of Machine Learning Research, 13(null):281–305.

Blundell, I., Brette, R., Cleland, T. A., Close, T. G., Coca, D., Davison, A. P., Diaz-Pier, S., Fernandez Musoles, C., Gleeson, P., Goodman, D. F. M., Hines, M., Hopkins, M. W., Kumbhar, P., Lester, D. R., Marin, B., Morrison, A., Müller, E., Nowotny, T., Peyser, A., Plotnikov, D., Richmond, P., Rowley, A., Rumpe, B., Stimberg, M., Stokes, A. B., Tomkins, A., Trensch, G., Woodman, M., and Eppler, J. M. (2018). Code generation in computational neuroscience: A review of tools and techniques. Frontiers in Neuroinformatics, 12.

Boucsein, C., Nawrot, M. P., Schnepel, P., and Aertsen, A. (2011). Beyond the Cortical Column: Abundance and Physiology of Horizontal Connections Imply a Strong Role for Inputs from the Surround. Frontiers in Neuroscience, 5.

Braitenberg, V. and Schüz, A. (1998). Cortex: Statistics and Geometry of Neuronal Connectivity. Springer Berlin Heidelberg, Berlin, Heidelberg.

Brette, R. and Goodman, D. F. (2012). Simulating spiking neural networks on GPU. Network: Computation in Neural Systems, 23(4):167–182.

Brette, R., Rudolph, M., Carnevale, T., Hines, M., Beeman, D., Bower, J. M., Diesmann, M., Morrison, A., Goodman, P. H., Harris, F. C., Zirpe, M., Natschläger, T., Pecevski, D., Ermentrout, B., Djurfeldt, M., Lansner, A., Rochel, O., Vieville, T., Muller, E., Davison, A. P., El Boustani, S., and Destexhe, A. (2007). Simulation of networks of spiking neurons: A review of tools and strategies. Journal of Computational Neuroscience, 23(3):349–398.

Brunel, N. (2000). Dynamics of sparsely connected networks of excitatory and inhibitory spiking neurons. Journal of computational neuroscience, 8(3):183–208.

Bruzzone, M., Chiarello, E., Albanesi, M., Miletto Petrazzini, M. E., Megighian, A., Lodovichi, C., and Dal Maschio, M. (2021). Whole brain functional recordings at cellular resolution in zebrafish larvae with 3d scanning multiphoton microscopy. Scientific Reports, 11.

Büsing, L., Schrauwen, B., and Legenstein, R. (2010). Connectivity, dynamics, and memory in reservoir computing with binary and analog neurons. Neural computation, 22(5):1272–1311.

Carlson, K. D., Nageswaran, J. M., Dutt, N., and Krichmar, J. L. (2014). An efficient automated parameter tuning framework for spiking neural networks. Frontiers in Neuro-science, 8.

Chicca, E., Stefanini, F., Bartolozzi, C., and Indiveri, G. (2014). Neuromorphic Electronic Circuits for Building Autonomous Cognitive Systems. Proceedings of the IEEE, 102(9):1367–1388.

Deepu, R., Spreizer, S., Trensch, G., Terhorst, D., Vennemo, S. B., Mitchell, J., Linssen, C., Mørk, H., Morrison, A., Eppler, J. M., Kamiji, N. L., de Schepper, R., Kitayama, I., Kurth, A., Morales-Gregorio, A., Nagendra Babu, P., and Plesser, H. E. (2021). Nest 3.1. Zenodo.

Deger, M., Helias, M., Rotter, S., and Diesmann, M. (2012). Spike-timing dependence of structural plasticity explains cooperative synapse formation in the neocortex. PLOS Computational Biology, 8(9):1–13.

Diesmann, M. and Gewaltig, M.-O. (2002). NEST: An Environment for Neural Systems. Forschung und Wisschenschaftliches Rechnen Beiträge zum Heinz-Billing-Preis, 58.

Diesmann, M., Gewaltig, M.-O., and Aertsen, A. (1995). Synod: An environment for neural systems simulations language interface and tutorial. Technical report, The Weizmann Institute of Science, 76100 Rehovot.

Diesmann, M., Gewaltig, M.-O., and Aertsen, A. (1999). Stable propagation of synchronous spiking in cortical neural networks. Nature, 402(6761):529–533.

Eliasmith, C. and Anderson, C. H. (2003). Neural engineering: Computation, representation, and dynamics in neurobiological systems. MIT press.

Eliasmith, C., Stewart, T. C., Choo, X., Bekolay, T., DeWolf, T., Tang, Y., and Rasmussen, D. (2012). A large-scale model of the functioning brain. Science, 338(6111):1202–1205.

Eliasmith, C. and Trujillo, O. (2014). The use and abuse of large-scale brain models. Current Opinion in Neurobiology, 25:1–6.

Eppler, J. M., Helias, M., Muller, E., Diesmann, M., and Gewaltig, M.-O. (2009). PyNEST: a convenient interface to the NEST simulator. Frontiers in Neuroinformatics, 2.

Eschbach, C. and Zlatic, M. (2020). Useful road maps: studying Drosophila larva’s central nervous system with the help of connectomics. Current Opinion in Neurobiology, 65:129–137.

Feldotto, B., Eppler, J. M., Jimenez-Romero, C., Bignamini, C., Gutierrez, C. E., Albanese, U., Retamino, E., Vorobev, V., Zolfaghari, V., Upton, A., Sun, Z., Yamaura, H., Heidarinejad, M., Klijn, W., Morrison, A., Cruz, F., McMurtrie, C., Knoll, A. C., Igarashi, J., Yamazaki, T., Doya, K., and Morin, F. O. (2022). Deploying and Optimizing Embodied Simulations of Large-Scale Spiking Neural Networks on HPC Infrastructure. Frontiers in Neuroinformatics, 16:884180.

Feurer, M. and Hutter, F. (2019). Hyperparameter Optimization, pages 3–33. Springer International Publishing, Cham.

Fidjeland, A. K., Roesch, E. B., Shanahan, M. P., and Luk, W. (2009). NeMo: a platform for neural modelling of spiking neurons using GPUs. In 2009 20th IEEE international conference on application-specific systems, architectures and processors, pages 137–144. IEEE.

Finkelstein, A., Fontolan, L., Economo, M. N., Li, N., Romani, S., and Svoboda, K. (2021). Attractor dynamics gate cortical information flow during decision-making. Nature Neuroscience, 24(6):843–850.

Florimbi, G., Torti, E., Masoli, S., D’Angelo, E., and Leporati, F. (2021). Granular layEr Simulator: Design and Multi-GPU Simulation of the Cerebellar Granular Layer. Frontiers in Computational Neuroscience, 15.

Gallinaro, J. V., Gašparović, N., and Rotter, S. (2022). Homeostatic control of synaptic rewiring in recurrent networks induces the formation of stable memory engrams. PLOS Computational Biology, 18(2):1–40.

Gewaltig, M.-O. and Diesmann, M. (2007). Nest (neural simulation tool). Scholarpedia, 2(4):1430.

Gjorgjieva, J., Drion, G., and Marder, E. (2016). Computational implications of biophysical diversity and multiple timescales in neurons and synapses for circuit performance. Current Opinion in Neurobiology, 37:44–52.

Godfrey, R. K., Swartzlander, M., and Gronenberg, W. (2021). Allometric analysis of brain cell number in Hymenoptera suggests ant brains diverge from general trends. Proceedings of the Royal Society B: Biological Sciences, 288(1947):20210199.

Golosio, B., Tiddia, G., De Luca, C., Pastorelli, E., Simula, F., and Paolucci, P. S. (2021). Fast simulations of highly-connected spiking cortical models using gpus. Frontiers in Computational Neuroscience, 15.

Gütig, R. (2016). Spiking neurons can discover predictive features by aggregate-label learning. Science, 351(6277):aab4113.

Gütig, R. and Sompolinsky, H. (2006). The tempotron: a neuron that learns spike timing– based decisions. Nature neuroscience, 9(3):420–428.

Hammond, C., Bergman, H., and Brown, P. (2007). Pathological synchronization in Parkinson’s disease: networks, models and treatments. Trends in neurosciences, 30(7):357–364.

Hazan, H., Saunders, D. J., Khan, H., Patel, D., Sanghavi, D. T., Siegelmann, H. T., and Kozma, R. (2018). BindsNET: A Machine Learning-Oriented Spiking Neural Networks Library in Python. Frontiers in Neuroinformatics, 12:89.

Helgadóttir, L. I., Haenicke, J., Landgraf, T., Rojas, R., and Nawrot, M. P. (2013). Conditioned behavior in a robot controlled by a spiking neural network. In 2013 6th International IEEE/EMBS Conference on Neural Engineering (NER), pages 891–894.

Herculano-Houzel, S., Watson, C., and Paxinos, G. (2013). Distribution of neurons in functional areas of the mouse cerebral cortex reveals quantitatively different cortical zones. Frontiers in neuroanatomy, 7:35.

Hines, M. L. and Carnevale, N. T. (2001). NEURON: a tool for neuroscientists. The neuroscientist, 7(2):123–135.

Hinsch, K. and Zupanc, G. (2007). Generation and long-term persistence of new neurons in the adult zebrafish brain: A quantitative analysis. Neuroscience, 146(2):679–696.

Hopfield, J. J. (1982). Neural networks and physical systems with emergent collective computational abilities. Proceedings of the National Academy of Sciences of the United States of America, 79(8):2554–2558.

Igarashi, J., Yamaura, H., and Yamazaki, T. (2019). Large-Scale Simulation of a Layered Cortical Sheet of Spiking Network Model Using a Tile Partitioning Method. Frontiers in Neuroinformatics, 13.

Illing, B., Ventura, J., Bellec, G., and Gerstner, W. (2021). Local plasticity rules can learn deep representations using self-supervised contrastive predictions. In Ranzato, M., Beygelzimer, A., Dauphin, Y., Liang, P., and Vaughan, J. W., editors, Advances in Neural Information Processing Systems, volume 34, pages 30365–30379. Curran Associates, Inc.

Inagaki, H. K., Fontolan, L., Romani, S., and Svoboda, K. (2019). Discrete attractor dynamics underlies persistent activity in the frontal cortex. Nature, 566(7743):212–217.

Indiveri, G., Stefanini, F., and Chicca, E. (2010). Spike-based learning with a generalized integrate and fire silicon neuron. In Proceedings of 2010 IEEE International Symposium on Circuits and Systems, \xpages 1951–1954.

Ippen, T., Eppler, J. M., Plesser, H. E., and Diesmann, M. (2017). Constructing neuronal network models in massively parallel environments. Frontiers in Neuroinformatics, 11.

Ivanov, D., Chezhegov, A., Grunin, A., Kiselev, M., and Larionov, D. (2022). Neuromorphic Artificial Intelligence Systems.

Javanshir, A., Nguyen, T. T., Mahmud, M. A. P., and Kouzani, A. Z. (2022). Advancements in Algorithms and Neuromorphic Hardware for Spiking Neural Networks. Neural Computation, 34(6):1289–1328.

Jordan, J., Ippen, T., Helias, M., Kitayama, I., Sato, M., Igarashi, J., Diesmann, M., and Kunkel, S. (2018). Extremely Scalable Spiking Neuronal Network Simulation Code: From Laptops to Exascale Computers. Frontiers in Neuroinformatics, 12:2.

Kasabov, N. and Capecci, E. (2015). Spiking neural network methodology for modelling, classification and understanding of EEG spatio-temporal data measuring cognitive processes. Information Sciences, 294:565–575.

Knight, J. C., Komissarov, A., and Nowotny, T. (2021). PyGeNN: A Python Library for GPU-Enhanced Neural Networks. Frontiers in Neuroinformatics, 15.

Knight, J. C. and Nowotny, T. (2018). GPUs Outperform Current HPC and Neuromorphic Solutions in Terms of Speed and Energy When Simulating a Highly-Connected Cortical Model. Frontiers in Neuroscience, 12.

Knight, J. C. and Nowotny, T. (2021). Larger GPU-accelerated brain simulations with procedural connectivity. Nature Computational Science, 1(2):136–142.

Knight, J. C. and Nowotny, T. (2022). Efficient GPU Training of LSNNs Using EProp. In Neuro-Inspired Computational Elements Conference, NICE 2022, page 8–10, New York, NY, USA. Association for Computing Machinery.

Kulesza, R. J. (2007). Cytoarchitecture of the human superior olivary complex: Medial and lateral superior olive. Hearing Research, 225(1):80–90.

Kulkarni, S. R., Parsa, M., Mitchell, J. P., and Schuman, C. D. (2021). Benchmarking the performance of neuromorphic and spiking neural network simulators. Neurocomputing, 447:145–160.

Kunkel, S., Potjans, T., Eppler, J., Plesser, H. E., Morrison, A., and Diesmann, M. (2012). Meeting the Memory Challenges of Brain-Scale Network Simulation. Frontiers in Neu-roinformatics, 5.

Kunkel, S., Schmidt, M., Eppler, J. M., Plesser, H. E., Masumoto, G., Igarashi, J., Ishii, S., Fukai, T., Morrison, A., Diesmann, M., and Helias, M. (2014). Spiking network simulation code for petascale computers. Frontiers in Neuroinformatics, 8.

Kurth, A. C., Senk, J., Terhorst, D., Finnerty, J., and Diesmann, M. (2022). Sub-realtime simulation of a neuronal network of natural density. Neuromorphic Computing and Engineering, 2(2):021001.

LaValle, S. M., Branicky, M. S., and Lindemann, S. R. (2004). On the Relationship between Classical Grid Search and Probabilistic Roadmaps. The International Journal of Robotics Research, (7-8):673–692.

Lengler, J., Jug, F., and Steger, A. (2013). Reliable Neuronal Systems: The Importance of Heterogeneity. PLoS ONE, 8(12):e80694.

Litwin-Kumar, A. and Doiron, B. (2012). Slow dynamics and high variability in balanced cortical networks with clustered connections. Nature Neuroscience, 15(11):1498–1505.

Litwin-Kumar, A., Rosenbaum, R., and Doiron, B. (2016). Inhibitory stabilization and visual coding in cortical circuits with multiple interneuron subtypes. Journal of Neurophysiology, 115(3):1399–1409.

Markram, H., Muller, E., Ramaswamy, S., Reimann, M. W., Abdellah, M., Sanchez, C. A., Ailamaki, A., Alonso-Nanclares, L., Antille, N., Arsever, S., Kahou, G. A. A., Berger, T. K., Bilgili, A., Buncic, N., Chalimourda, A., Chindemi, G., Courcol, J.-D., Delalondre, F., Delattre, V., Druckmann, S., Dumusc, R., Dynes, J., Eilemann, S., Gal, E., Gevaert, M. E., Ghobril, J.-P., Gidon, A., Graham, J. W., Gupta, A., Haenel, V., Hay, E., Heinis, T., Her-nando, J. B., Hines, M., Kanari, L., Keller, D., Kenyon, J., Khazen, G., Kim, Y., King, J. G., Kisvarday, Z., Kumbhar, P., Lasserre, S., Le Bé, J.-V., Magalhães, B. R., Merchán-Pérez, A., Meystre, J., Morrice, B. R., Muller, J., Muñoz-Céspedes, A., Muralidhar, S., Muthurasa, K., Nachbaur, D., Newton, T. H., Nolte, M., Ovcharenko, A., Palacios, J., Pastor, L., Perin, R., Ranjan, R., Riachi, I., Rodríguez, J.-R., Riquelme, J. L., Rössert, C., Sfyrakis, K., Shi, Y., Shillcock, J. C., Silberberg, G., Silva, R., Tauheed, F., Telefont, M., Toledo-Rodriguez, M., Tränkler, T., Van Geit, W., Díaz, J. V., Walker, R., Wang, Y., Zaninetta, S. M., DeFelipe, J., Hill, S. L., Segev, I., and Schürmann, F. (2015). Reconstruction and simulation of neocortical microcircuitry. Cell, 163(2):456–492.

Mazzucato, L. (2022). Neural mechanisms underlying the temporal organization of naturalistic animal behavior. arXiv.

Mazzucato, L., Fontanini, A., and La Camera, G. (2015). Dynamics of multistable states during ongoing and evoked cortical activity. The Journal of neuroscience, 35(21):8214–31.

Mazzucato, L., La Camera, G., and Fontanini, A. (2019). Expectation-induced modulation of metastable activity underlies faster coding of sensory stimuli. Nature Neuroscience, 22:787–796.

McIntyre, C. C. and Hahn, P. J. (2010). Network perspectives on the mechanisms of deep brain stimulation. Neurobiology of disease, 38(3):329–337.

Mejias, J. F. and Longtin, A. (2012). Optimal Heterogeneity for Coding in Spiking Neural Networks. Physical Review Letters, 108(22):228102.

Mejias, J. F. and Longtin, A. (2014). Differential effects of excitatory and inhibitory heterogeneity on the gain and asynchronous state of sparse cortical networks. Frontiers in Computational Neuroscience, 8.

Menzel, R. (2012). The honeybee as a model for understanding the basis of cognition. Nature reviews. Neuroscience, 13:758–68.

Morrison, A., Mehring, C., Geisel, T., Aertsen, A., and Diesmann, M. (2005). Advancing the Boundaries of High-Connectivity Network Simulation with Distributed Computing. Neural Computation, 17(8):1776–1801.

Morrison, A., Straube, S., Plesser, H. E., and Diesmann, M. (2007). Exact subthreshold integration with continuous spike times in discrete-time neural network simulations. Neural computation, 19(1):47–79.

Mutch, J., Knoblich, U., and Poggio, T. (2010). CNS: a GPU-based framework for simulating cortically-organized networks. MIT CSAIL, 17:2013.

Nageswaran, J. M., Dutt, N., Krichmar, J. L., Nicolau, A., and Veidenbaum, A. V. (2009). A configurable simulation environment for the efficient simulation of large-scale spiking neural networks on graphics processors. Neural Networks, 22(5):791–800.

Nawrot, M. P., Schnepel, P., Aertsen, A., and Boucsein, C. (2009). Precisely timed signal transmission in neocortical networks with reliable intermediate-range projections. Frontiers in Neural Circuits, 3.

Neftci, E., Binas, J., Rutishauser, U., Chicca, E., Indiveri, G., and Douglas, R. J. (2013). Synthesizing cognition in neuromorphic electronic systems. Proceedings of the National Academy of Sciences of the United States of America, 110(37).

Niedermeier, L., Chen, K., Xing, J., Das, A., Kopsick, J., Scott, E., Sutton, N., Weber, K., Dutt, N., and Krichmar, J. L. (2022). CARLsim 6: An Open Source Library for Large-Scale, Biologically Detailed Spiking Neural Network Simulation. In 2022 International Joint Conference on Neural Networks (IJCNN), pages 1–10, Padua, Italy. IEEE.

Oorschot, D. E. (1996). Total number of neurons in the neostriatal, pallidal, subthalamic, and substantia nigral nuclei of the rat basal ganglia: A stereological study using the cavalieri and optical disector methods. Journal of Comparative Neurology, 366(4):580–599.

Ostrau, C., Klarhorst, C., Thies, M., and Rückert, U. (2022). Benchmarking Neuromorphic Hardware and Its Energy Expenditure. Frontiers in Neuroscience, 16:873935.

Parrish-Aungst, S., Shipley, M., Erdelyi, F., Szabó, G., and Puche, A. (2007). Quantitative analysis of neuronal diversity in the mouse olfactory bulb. The Journal of comparative neurology, 501:825–36.

Parsa, M., Mitchell, J. P., Schuman, C. D., Patton, R. M., Potok, T. E., and Roy, K. (2019). Bayesian-based Hyperparameter Optimization for Spiking Neuromorphic Systems. In 2019 IEEE International Conference on Big Data (Big Data), pages 4472–4478.

Pfeiffer, M. and Pfeil, T. (2018). Deep Learning With Spiking Neurons: Opportunities and Challenges. Frontiers in Neuroscience, 12.

Plotnikov, D., Rumpe, B., Blundell, I., Ippen, T., Eppler, J. M., and Morrison, A. (2016). NESTML: a modeling language for spiking neurons. In Modellierung 2016, March 2–4 2016.

Potjans, T. C. and Diesmann, M. (2014). The Cell-Type Specific Cortical Microcircuit: Relating Structure and Activity in a Full-Scale Spiking Network Model. Cerebral Cortex, 24(3):785–806.

Pronold, J., Jordan, J., Wylie, B. J. N., Kitayama, I., Diesmann, M., and Kunkel, S. (2022). Routing brain traffic through the von Neumann bottleneck: Efficient cache usage in spiking neural network simulation code on general purpose computers. Parallel Computing, 113:102952.

Raji, J. I. and Potter, C. J. (2021). The number of neurons in Drosophila and mosquito brains. PLOS ONE, 16(5):1–11.

Rapp, H. and Nawrot, M. P. (2020). A spiking neural program for sensorimotor control during foraging in flying insects. Proceedings of the National Academy of Sciences, 117(45):28412–28421.

Rapp, H., Nawrot, M. P., and Stern, M. (2020). Numerical Cognition Based on Precise Counting with a Single Spiking Neuron. iScience, 23(2):100852.

Rost, T. (2016). Modelling Cortical Variability Dynamics. PhD thesis, Freie Universität Berlin.

Rost, T., Deger, M., and Nawrot, M. P. (2018). Winnerless competition in clustered balanced networks: inhibitory assemblies do the trick. Biological cybernetics, 112(1-2):81–98.

Rostami, V., Rost, T., Riehle, A., van Albada, S. J., and Nawrot, M. P. (2022). Excitatory and inhibitory motor cortical clusters account for balance, variability, and task performance. bioRxiv.

Rotter, S. and Diesmann, M. (1999). Exact digital simulation of time-invariant linear systems with applications to neuronal modeling. Biological Cybernetics, 81(5):381–402.

Sacramento, J. a., Ponte Costa, R., Bengio, Y., and Senn, W. (2018). Dendritic cortical microcircuits approximate the backpropagation algorithm. In Bengio, S., Wallach, H., Larochelle, H., Grauman, K., Cesa-Bianchi, N., and Garnett, R., editors, Advances in Neural Information Processing Systems, volume 31. Curran Associates, Inc.

Sakagiannis, P., Jürgensen, A.-M., and Nawrot, M. P. (2021). A realistic locomotory model of drosophila larva for behavioral simulations. bioRxiv.

Sakai, K. and Miyashita, Y. (1991). Neural organization for the long-term memory of paired associates. Nature, 354(6349):152–155.

Sarko, D. K., Catania, K., Leitch, D. B., Kaas, J. H., and Herculano-Houzel, S. (2009). Cellular scaling rules of insectivore brains. Frontiers in Neuroanatomy, 3.

Schmidt, M., Bakker, R., Shen, K., Bezgin, G., Diesmann, M., and van Albada, S. J. (2018). A multi-scale layer-resolved spiking network model of resting-state dynamics in macaque visual cortical areas. PLOS Computational Biology, 14(10):1–38.

Schmuker, M., Pfeil, T., and Nawrot, M. P. (2014). A neuromorphic network for generic multivariate data classification. Proceedings of the National Academy of Sciences, 111(6):2081–2086.

Schuman, C. D., Kulkarni, S. R., Parsa, M., Mitchell, J. P., Date, P., and Kay, B. (2022). Opportunities for neuromorphic computing algorithms and applications. Nature Computational Science, 2(1):10–19.

Sherwood, C. C., Miller, S. B., Karl, M., Stimpson, C. D., Phillips, K. A., Jacobs, B., Hof, P. R., Raghanti, M. A., and Smaers, J. B. (2020). Invariant Synapse Density and Neuronal Connectivity Scaling in Primate Neocortical Evolution. Cerebral Cortex, 30(10):5604–5615.

Singer, W. and Gray, C. M. (1995). Visual feature integration and the temporal correlation hypothesis. Annual review of neuroscience, 18(1):555–586.

Steffen, L., Koch, R., Ulbrich, S., Nitzsche, S., Roennau, A., and Dillmann, R. (2021). Benchmarking Highly Parallel Hardware for Spiking Neural Networks in Robotics. Frontiers in Neuroscience, 15.

Stimberg, M., Brette, R., and Goodman, D. F. M. (2019). Brian 2, an intuitive and efficient neural simulator. eLife, 8.

Tanaka, G., Yamane, T., Héroux, J. B., Nakane, R., Kanazawa, N., Takeda, S., Numata, H., Nakano, D., and Hirose, A. (2019). Recent advances in physical reservoir computing: A review. Neural Networks, 115:100–123.

Tavanaei, A., Ghodrati, M., Kheradpisheh, S. R., Masquelier, T., and Maida, A. (2019). Deep learning in spiking neural networks. Neural Networks, 111:47–63.

Thain, D., Tannenbaum, T., and Livny, M. (2005). Distributed computing in practice: the Condor experience. Concurrency - Practice and Experience, 17(2-4):323–356.

Thörnig, P. (2021). JURECA: Data Centric and Booster Modules implementing the Modular Supercomputing Architecture at Jülich Supercomputing Centre. Journal of large-scale research facilities JLSRF, 7:A182.

Tiddia, G., Golosio, B., Albers, J., Senk, J., Simula, F., Pronold, J., Fanti, V., Pastorelli, E., Paolucci, P. S., and van Albada, S. J. (2022). Fast Simulation of a Multi-Area Spiking Network Model of Macaque Cortex on an MPI-GPU Cluster. Frontiers in Neuroinformatics, 16:883333.

Tikidji-Hamburyan, R. A., Narayana, V., Bozkus, Z., and El-Ghazawi, T. A. (2017). Software for Brain Network Simulations: A Comparative Study. Frontiers in Neuroinformatics, 11.

Tripathy, S. J., Padmanabhan, K., Gerkin, R. C., and Urban, N. N. (2013). Intermediate intrinsic diversity enhances neural population coding. Proceedings of the National Academy of Sciences, 110(20):8248–8253.

Van Albada, S. J., Helias, M., and Diesmann, M. (2015). Scalability of asynchronous networks is limited by one-to-one mapping between effective connectivity and correlations. PLoS computational biology, 11(9):e1004490.

Van Vreeswijk, C. and Sompolinsky, H. (1996). Chaos in neuronal networks with balanced excitatory and inhibitory activity. Science, 274(5293):1724–1726.

Vitay, J., Dinkelbach, H. O., and Hamker, F. H. (2015). ANNarchy: a code generation approach to neural simulations on parallel hardware. Frontiers in Neuroinformatics, 9.

Vlag, M. A. v. d., Smaragdos, G., Al-Ars, Z., and Strydis, C. (2019). Exploring Complex Brain-Simulation Workloads on Multi-GPU Deployments. ACM Transactions on Architecture and Code Optimization, 16(4):1–25.

Vogels, T. P., Sprekeler, H., Zenke, F., Clopath, C., and Gerstner, W. (2011). Inhibitory plasticity balances excitation and inhibition in sensory pathways and memory networks. Science, 334(6062):1569–1573.

Von St. Vieth, B. (2021). JUSUF: Modular Tier-2 Supercomputing and Cloud Infrastructure at Jülich Supercomputing Centre. Journal of large-scale research facilities JLSRF, 7:A179.

White, J. G., Southgate, E., Thomson, J. N., and Brenner, S. (1986). The structure of the nervous system of the nematode Caenorhabditis elegans. Philosophical Transactions of the Royal Society of London. B, Biological Sciences, 314(1165):1–340.

Witthöft, W. (1967). Absolute Anzahl und Verteilung der Zellen im Hirn der Honigbiene. Zeitschrift für Morphologie der Tiere, 61(1):160–184.

Wyrick, D. and Mazzucato, L. (2021). State-dependent regulation of cortical processing speed via gain modulation. Journal of Neuroscience, 41(18):3988–4005.

Yamaura, H., Igarashi, J., and Yamazaki, T. (2020). Simulation of a Human-Scale Cerebellar Network Model on the K Computer. Frontiers in Neuroinformatics, 14.

Yavuz, E., Turner, J., and Nowotny, T. (2016). GeNN: a code generation framework for accelerated brain simulations. Scientific reports, 6(1):1–14.

Yegenoglu, A., Subramoney, A., Hater, T., Jimenez-Romero, C., Klijn, W., Pérez Martín, A., van der Vlag, M., Herty, M., Morrison, A., and Diaz-Pier, S. (2022). Exploring Parameter and Hyper-Parameter Spaces of Neuroscience Models on High Performance Computers With Learning to Learn. Frontiers in Computational Neuroscience, 16.

Zenke, F. and Ganguli, S. (2018). SuperSpike: Supervised Learning in Multilayer Spiking Neural Networks. Neural Computation, 30(6):1514–1541.

Zhao, Y. and Wang, Dan O. and Martin, K. C. (2009). Preparation of Aplysia sensory-motor neuronal cell cultures. Journal of visualized experiments : JoVE.

